# Domain Architecture Analysis Guides the Discovery of Unusual Glycyl Radical Enzyme Fusion Proteins

**DOI:** 10.64898/2026.11.23.768175

**Authors:** Gil Namkoong, Nina M. Greeley, Catherine L. Drennan, Emily P. Balskus

**Affiliations:** Department of Chemistry and Chemical Biology, Harvard University, Cambridge, MA 02138, USA; Department of Chemistry, Massachusetts Institute of Technology, Cambridge, MA 02139, USA; Department of Biology, Massachusetts Institute of Technology, Cambridge, MA 02139, USA; Howard Hughes Medical Institute, Massachusetts Institute of Technology, Cambridge, MA 02139, USA; Howard Hughes Medical Institute, Harvard University, Cambridge, MA 02138, USA

**Keywords:** radical chemistry, anaerobic metabolism, sequence similarity network, domain fusion, diatom, cryo-EM

## Abstract

Glycyl radical enzymes (GREs) use a post-translationally installed glycyl radical cofactor to catalyze chemically challenging reactions that are critical for anaerobic metabolism in microorganisms. Despite its importance, the GRE superfamily remains undercharacterized in part due to the vastness of its sequence space. Herein, we use sequence similarity network (SSN) analysis to survey the sequence diversity of the GRE superfamily and identify previously uncharacterized subfamilies. By analyzing the conserved domain architecture of each uncharacterized cluster, we find an unexpected cluster of GRE sequences from diverse photosynthetic eukaryotes. Bioinformatic, biochemical, and structural studies together demonstrate that these GREs have their catalytic subunits fused with a GRE-activating enzyme (GRE-AE). We characterize a highly expressed member of this subfamily as a functional pyruvate formate-lyase that is activated by its fused AE. Low resolution cryogenic electron microscopy confirms that this protein is a naturally occurring fusion, with a central dimeric GRE domain and one GRE-AE domain on each side. We further find that the GRE/GRE-AE fusion proteins are transcriptionally active in a wide range of diatoms and related microalgae. Overall, these findings uncover a previously unappreciated role for GRE-mediated anaerobic metabolism in ecologically important phototrophs, reveal that an attached AE can install a glycyl radical species on its cognate GRE, and show that domain architecture analysis is a powerful strategy to find novel enzymes among large enzyme superfamilies.

**Significance Statement:** Glycyl radical enzymes (GREs) mediate chemically challenging and physiologically important reactions in anaerobic microorganisms, yet only a fraction have known functions. Here, we perform sequence similarity network analysis of the GRE superfamily to identify uncharacterized GREs with atypical features. By targeting sequences with unusual combinations of domains for further study, we uncover a group of GREs that are natively fused to their partner GRE-activating enzymes (GRE-AEs). Our bioinformatic, biochemical, and structural studies show that GRE/GRE-AE fusion proteins are self-activating, functional pyruvate formate-lyases expressed in diverse microalgae that typically grow aerobically. Our findings expand the distribution of GRE-mediated metabolism beyond anaerobes and demonstrate the utility of domain architecture analysis for identifying novel enzymes in large superfamilies.

## Introduction

Glycyl radical enzymes (GREs) are evolutionarily ancient enzymes that play important roles in diverse anaerobic microbes (1–4). They use a post-translationally installed glycyl radical cofactor to perform chemically challenging transformations that are central to anaerobic microbial metabolism. This protein-based radical is installed at the α-carbon atom of a conserved glycine residue by the action of a GRE-activating enzyme (GRE-AE) that is typically specific to each GRE and is encoded by a neighboring gene to the gene that encodes the GRE (**Figure 1A**). GRE-AEs belong to the radical *S*-adenosyl-L-methionine (RS) enzyme superfamily, which reductively cleaves *S*-adenosyl-L-methionine (AdoMet) at the conserved [Fe_4_S_4_] cluster to generate a 5′-deoxyadenosyl radical (5′dAdo•) that abstracts a hydrogen atom (H-atom) from the conserved glycine of the cognate GRE (**Figure 1B**) (5, 6). This conserved glycine is located on a buried loop within the glycyl radical domain (GRD) that sits inside the enzyme’s active site pocket. It is believed that the interaction with GRE-AE induces the GRD to flip out into the GRE-AE’s active site (7–9). However, the molecular basis for the high specificity of the interaction between GRE and GRE-AE and for the GRE conformational change induced by GRE-AE are currently unknown.

**Figure 1.**
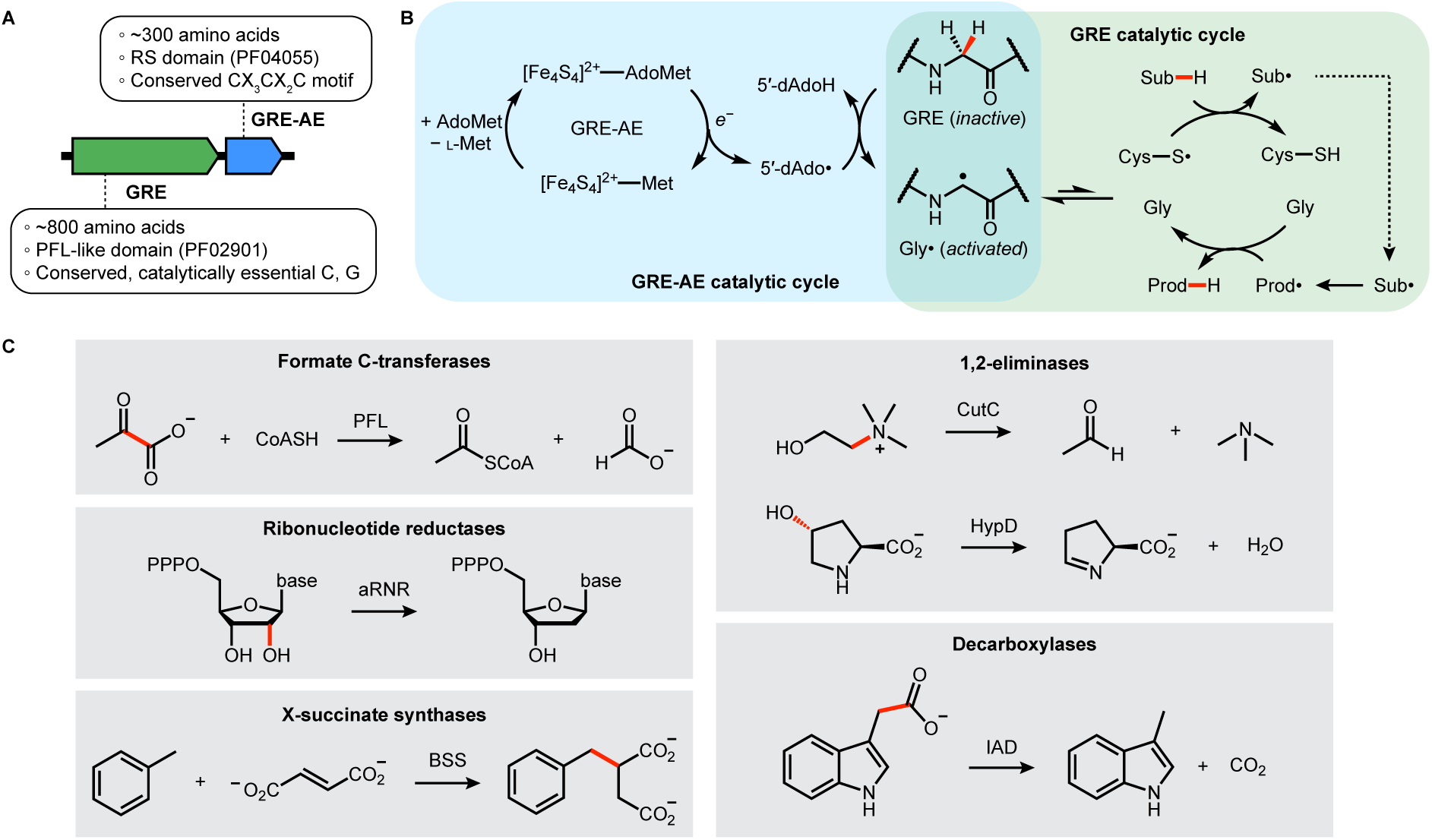
The glycyl radical enzyme (GRE) superfamily uses post-translationally installed protein-based radicals. **(A)** Typical organization of genes encoding a GRE and a GRE-activating enzyme (GRE-AE). RS = radical *S*-adenosyl-L-methionine. **(B)** General mechanism of GRE catalysis. The GRE is first activated by the GRE-AE (green box) before the activated glycyl radical is transferred to the conserved cysteine to initiate the radical chemistry on the substrate (purple box). AdoMet = *S*-adenosyl-L-methionine; 5′-dAdo• = 5′-deoxyadenosyl radical; 5′-dAdoH = 5′-deoxyadenosine; Sub = substrate; Prod = product. **(C)** Selected examples of GRE-mediated reactions. Bonds that are being broken or formed are highlighted in red. CoA = coenzyme A; PFL = pyruvate formate-lyase; PPP = triphosphate; aRNR = anaerobic (class III) ribonucleotide reductase; CutC = choline trimethylamine-lyase; HypD = *trans*-4-hydroxy-L-proline dehydratase; BSS = benzylsuccinate synthase; IAD = indole-3-acetate decarboxylase.

In the GRE catalytic cycle, the glycyl radical reacts with a conserved cysteine residue in the active site to yield a catalytically competent thiyl radical, which then typically abstracts an H-atom from the substrate to initiate the radical-based reaction (**Figure 1B**). There are ∼20 unique reactions currently known to be catalyzed by GREs that can be categorized into seven classes (**Figure 1C**): formate lyases, ribonucleotide reductases, 1,2-eliminases, isomerases, decarboxylases, X-succinate synthases (X = benzyl, naphthyl, or alkyl), and aminoacyl radical enzymes (3, 4). These reactions play important roles in microbial physiology and in host–microbe interactions. For example, pyruvate formate-lyase (PFL) is involved in anaerobic central metabolism (10, 11) and class III ribonucleotide reductase (RNR) is essential for DNA synthesis in anaerobes (12, 13). In addition, the activities of some GRE 1,2-eliminases and decarboxylases encoded by the human microbiome are associated with host health and disease states (14–17). Moreover, X-succinate synthases enable microbes in niche environments to utilize aromatic or aliphatic hydrocarbons as nutrients (18–20).

Despite their importance, the number of characterized GREs represents only a small fraction of the GRE superfamily (total >20k sequences in UniProtKB (21) in 2025). Herein, we leverage sequence similarity relationships to survey the sequence space of the GRE superfamily and identify unique subfamilies for functional characterization. By analyzing the domain architecture of each GRE cluster, we prioritize sequences with unusual domain fusions for biochemical characterization. Unexpectedly, we identify a novel cluster of GREs that are natively fused to their cognate GRE-AEs and are found in diatoms and closely related microalgae. *In vitro* biochemical characterization of a GRE/GRE-AE fusion protein from the diatom *Fistulifera solaris* demonstrates that the enzyme is a functional PFL. A structural reconstruction using cryogenic electron microscopy (cryo-EM) confirms that this protein is a fusion between a GRE and a GRE-AE. Bioinformatic analyses indicate that these GRE/GRE-AE fusion proteins are widely distributed across taxonomically and ecologically diverse diatoms and are highly expressed in these organisms. Altogether, this work reveals an unprecedented domain architecture for GREs, a previously unappreciated role for these enzymes in aerobic microbes, and the power of domain analysis in guiding enzyme discovery.

## Results

### Sequence similarity network (SSN) analysis of the GRE superfamily reveals novel, uncharacterized clusters

We previously utilized SSN analysis to identify and characterize new members of the GRE superfamily (16). As the number of total protein sequences in the UniProt database nearly tripled from ∼80 million to ∼240 million since our initial study, we sought to repeat the analysis to capture any new function that might be encoded in the current set of GRE sequences. We started our analysis by constructing a series of SSNs of the GRE superfamily (16, 22–25). Specifically, 20,907 sequences from UniProt database (version 2025-03) that contain PFL-like domain (Pfam 02901; does not include class III RNR) were submitted to the Enzyme Function Initiative Enzyme Similarity Tool (EFI-EST) for SSN generation (24, 25). We selected a series of alignment score (AS) threshold values to group these sequences into isofunctional clusters and to understand similarity relationships between different clusters. Similar to our previous study (16), an AS threshold of 200, which corresponds to an average amino acid sequence identity of 58%, effectively separated experimentally characterized GRE sequences into isofunctional clusters (**Figure 2A**, *left*). A few exceptions include: (i) cluster 7 containing both glycerol dehydratase (GD) (26) and propane-1,2-diol dehydratase (PD) (16, 27), which catalyze the same reaction on nearly identical substrates, (ii) cluster 9 that contains isethionate sulfite-lyase (IslA) (28, 29) and 2,3-dihydroxypropanesulfonate sulfolyase (HpsG) (30), both of which catalyze C–S bond cleavage on structurally similar substrates, and (iii) cluster 1 and cluster 5, which both contain PFL sequences with cluster 5 restricted to sequences from Streptococci and Lactococci. This SSN shows that although multiple GREs have been characterized over the past decade, the functions of many clusters remain unknown, reiterating the importance of creative approaches to prioritize the study of these uncharacterized enzymes.

**Figure 2.**
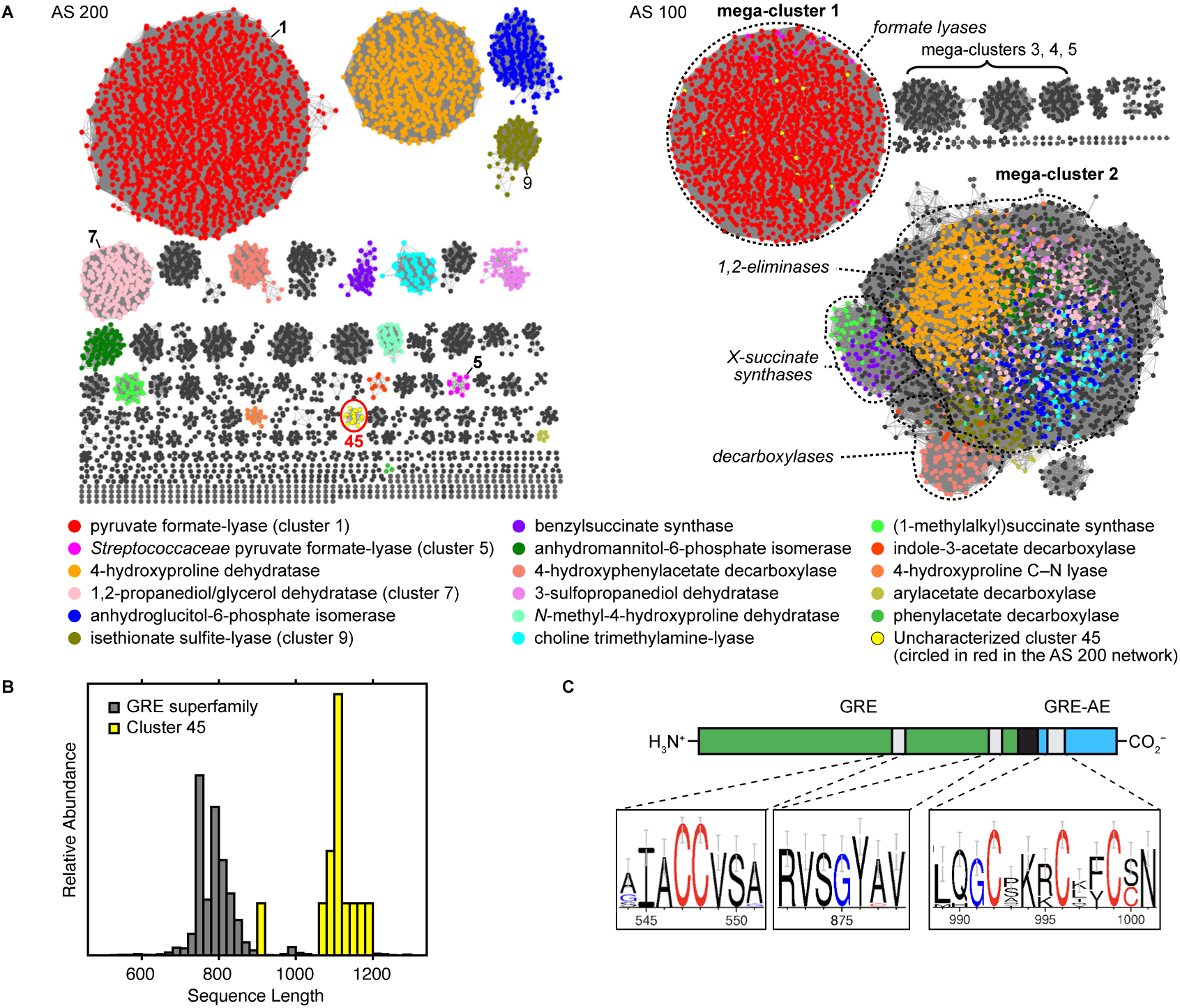
Sequence similarity network (SSN) analysis of the GRE superfamily (PF02901) identifies an unusual cluster consisting of GRE/GRE-AE fusion proteins. **(A)** SSNs of PF02901-containing sequences at alignment score threshold of 200 (*left*) and 100 (*right*). Each node represents 85% identity representative sequence. Clusters with characterized members are colored, whereas gray clusters are uncharacterized. The node colors used in the AS 200 network were used to color the corresponding nodes in the AS 100 network. Clusters referred to by number in the text are numbered on the panel. **(B)** Length histogram of Cluster 45 in comparison to the whole superfamily (85% representative sequences). **(C)** Conserved domain architecture of Cluster 45. Sequence logos of highly conserved regions are shown.

Starting from this SSN, we gradually decreased the threshold value to further evaluate the similarity relationships between the isofunctional clusters (**Figure S1–S2**) (22). As the AS threshold decreases, some clusters start to coalesce, indicating that they are more likely to be functionally related to each other. For example, at AS threshold of 150, the clusters corresponding to YbiW, HypD, PflD, PD, GD, and HpfG, all of which catalyze C– O bond cleavage reactions, merge to form a larger ‘dehydratase-type cluster’ that includes some 1,2-eliminases and isomerases (**Figure S1A** and **Figure S2**). At AS threshold of 125, the CutC, HpyG, HpzG, and IslA clusters also merge with the dehydratase-type cluster to form a ‘1,2-eliminase cluster’ (**Figure S1B** and **Figure S2**). Similarly, the HPAD and IAD clusters form a larger ‘decarboxylase cluster’ at this threshold, and the BSS and MASS clusters form a larger ‘XSS cluster’ at AS threshold of 120 (**Figure S1C–D** and **Figure S2**).

At an AS threshold of 100, more than 98% of all sequences fall into one of the five ‘mega-clusters’ (**Figure 2A**, *right*). Mega-cluster 1 contains clusters 1 and 5 from the AS 200 SSN—these clusters consist of PFLs and closely related TdcEs that catalyze formate lyase reactions. All other characterized GREs (excluding class III RNR) are found in mega-cluster 2, while different GRE classes are spatially segregated within this mega-cluster. It is notable that the reactivity of the thiyl radical of PFLs and all other characterized GREs are fundamentally different (**Figure S3**): thiyl radicals of GREs in mega-cluster 2 perform H-atom abstraction (or equivalent proton-coupled electron transfer) on substrates to generate a substrate-based radical species, whereas the thiyl radical of PFLs (mega-cluster 1) is first transferred to an adjacent, conserved cysteine residue, and this new thiyl radical performs a radical addition reaction to the substrate carbonyl group forming a covalent intermediate. Mega-clusters 3, 4, and 5 do not have any characterized members, implying that the GREs in these mega-clusters could perhaps display fundamentally distinct thiyl radical reactivities. This analysis provides a functional map of the GRE superfamily as a first step toward prioritizing uncharacterized clusters for investigation and predicting their reaction class.

### Cluster 45 contains GRE/GRE-AE fusion proteins from diatoms and other microalgae

Identification of novel enzymes from an SSN can be facilitated by mapping orthogonal information, such as known ligand/substrate, taxonomy, neighboring genes, or metagenomic abundance, onto individual clusters (16, 22). One strategy that has facilitated enzyme discovery in other superfamilies (22, 31, 32) but has not yet been utilized for the GRE superfamily is prioritizing sequences with unusual domain architectures. Observation of an unusual conserved fused domain architecture could also help to inform functional and/or mechanistic hypotheses. Cluster 45 stands out within the SSN (AS threshold 200) because it contains unusually long sequences compared to other sequences in the superfamily (1105 vs. 798 amino acids on average) (**Figure 2B**). Unexpectedly, analysis of the sequences in cluster 45 revealed the presence of Pfam 02901 (PFL-like domain) and Pfam 01228 (glycyl radical domain) in addition to Pfam 04055 (RS superfamily domain), suggesting that they could be fusions of a GRE and a GRE-AE. This domain architecture was reported in a previous study which identified a single putative fusion of PFL and PFL-AE in the diatom *Thalassiosira pseudonana* through a phylogenetic analysis of putative PFL sequences from eukaryotes (33). However, whether this fusion protein is a functionally active PFL and the prevalence of this fused domain architecture have not been explored.

Cluster 45 is a part of mega-cluster 1, indicating that its members are likely functionally related to PFLs. Consistent with this prediction, alignment of sequences from this cluster reveals the highly conserved ‘CCVS’ motif that is characteristic of PFLs (**Figure 2C**). These sequences also share a conserved Gly loop in the GRD that resembles the conserved Gly loop of *bona fide* PFLs (cluster 1). In the RS domain, the expected ‘CxxxCxxC’ motif for [Fe_4_S_4_] cluster binding is present, and there are no other conserved cysteine residues that could bind auxiliary FeS clusters. Altogether, the sequence analysis and the domain prediction suggest that these sequences are GRE/GRE-AE fusion proteins that likely function as PFLs.

In addition to its unusual domain architecture, cluster 45 is also notable for its taxonomic distribution. While more than 95% of sequences in the SSN originate from prokaryotes, sequences from cluster 45 are found exclusively in eukaryotes with the majority from *Bacillariophyta*, the phylum of diatoms. Diatoms are photosynthetic microalgae that inhabit various ecological niches and possess diverse metabolic capabilities. They are considered as aerobic organisms although they can survive under dark, anoxic conditions for some period using nitrate respiration (34). It was surprising to find a GRE cluster from diatoms, because the vast majority of GREs are found in facultative or obligate anaerobes, potentially due to their oxygen sensitivity. A few limited examples of previously identified eukaryotic GREs are PFLs from anaerobic fungi or amoeboid protists (33, 35–37). There is also a biochemically characterized PFL from the chloroplast and mitochondrion of a green alga *Chlamydomonas reinhardtii* (38, 39), a subcellular localization that could protect the enzyme from oxygen. In contrast, cluster 45 sequences do not contain signal peptides. All of these previously identified eukaryotic PFLs are found in cluster 1 along with other characterized bacterial PFLs and are not fused to a GRE-AE, making cluster 45 sequences distinct from other eukaryotic PFLs.

### Heterologously expressed *F. solaris* GRE/GRE-AE harbors an [Fe_4_S_4_] cluster and a glycyl radical

To validate the function of GRE/GRE-AE fusion proteins, a GRE/GRE-AE from *Fistulifera solaris* (UniProt ID A0A1Z5KPM5) was heterologously expressed in *Escherichia coli*. This protein was chosen as the representative member because it is highly expressed in the native organism (*vide infra*), has favorable predicted physical properties (e.g., isoelectric point, stability index), and is from a genetically tractable organism (40). Micro-aerobically expressed and anaerobically purified *Fs*GRE/GRE-AE is brown, contains *ca.* 4 Fe per polypeptide (**Table S1**), and features a characteristic absorption peak around 410 nm, indicative of an [Fe_4_S_4_] cluster (**Figure 3A**). The EPR spectrum of *Fs*GRE/GRE-AE reduced with sodium dithionite (NaDT) shows a rhombic *S* = 1/2 signal, consistent with the presence of an [Fe_4_S_4_]^+^ cluster (**Figure 3B**). This signal cleanly converts to a distinct *S* = ^1^/_2_ signal upon addition of AdoMet, indicating binding of AdoMet to the [Fe_4_S_4_]^+^ cluster as observed in *E. coli* PFL-AE (41) as well as other RS enzymes (6). To test whether the [Fe_4_S_4_] cluster is bound at the GRE-AE domain, we generated the C878A variant in which the first cysteine of the CxxxCxxC motif for [Fe_4_S_4_]-binding is mutated to an alanine (**Figure S4A**). The C878A variant is purified as a colorless protein lacking bound Fe, and its UV-Vis spectrum is consistent with the loss of the [Fe_4_S_4_] cluster (**Figure S4B**).

**Figure 3.**
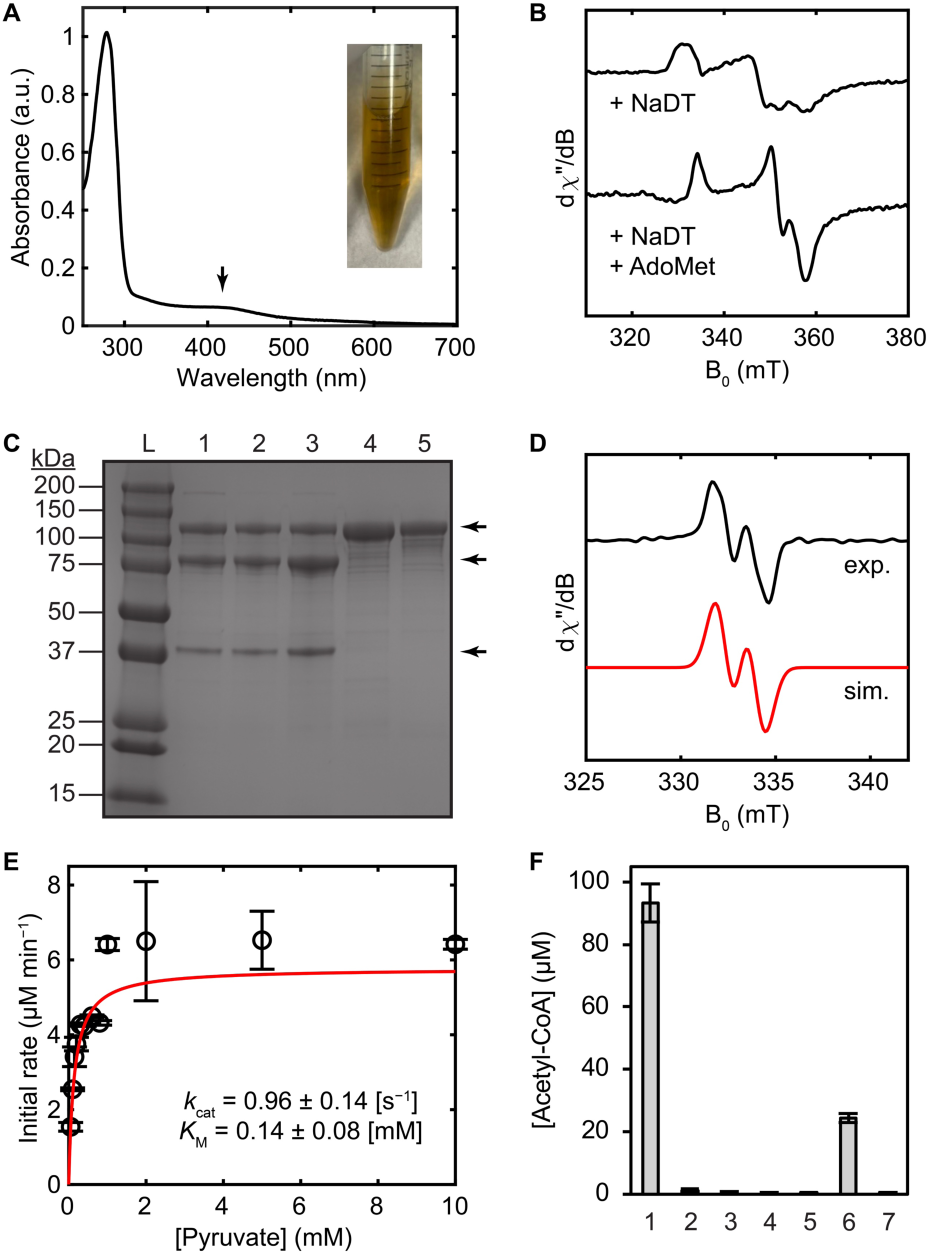
Biochemical characterization of a GRE/GRE-AE fusion protein from *Fistulifera solaris* shows that it is a functional PFL. **(A)** UV-Vis spectrum of as-isolated *Fs*GRE/GRE-AE fusion protein. The feature around 410 nm (marked with a black arrow) is indicative of a [Fe_4_S_4_]^2+^ cluster. (*Inset*) A photo of as-isolated *Fs*GRE/GRE-AE showing the characteristic brown color of a bound-FeS cluster. **(B)** 15 K EPR spectra of *Fs*GRE/GRE-AE in the presence of NaDT alone or with AdoMet. EPR conditions: frequency = 9.37 GHz; power = 0.25 mW (see SI for additional details). **(C)** SDS-PAGE of *Fs*GRE/GRE-AE variants. L: protein ladder; Lane 1: wild type (WT); Lane 2: C461S; Lane 3: C462S; Lane 4: G785A; Lane 5: C878A. Three bands corresponding to the full-length protein (126 kDa), *N*-terminal fragment (90 kDa), and *C*-terminal fragment (36 kDa) are marked with arrows. **(D)** 77 K EPR spectrum of as-isolated *Fs*GRE/GRE-AE (top black trace) and its simulation (bottom red trace). EPR conditions: 9.37 GHz; power = 1 µW (see SI for additional details). Simulation parameters: *g*_iso_ = 2.01; *g*-strain = 0.0051; **A**(^1^H) = [52.1, 28.5, 42.6] MHz; linewidth = 1.1 mT. **(E)** Michaelis–Menten kinetics of *Fs*GRE/GRE-AE (activated with Ti^III^-citrate) under varying concentrations of pyruvate. Each data point with an error bar represents mean ± S.D. (standard deviation) of duplicate measurements. Assay conditions: [Enzyme] = 0.1 µM; [CoA] = 0.1 mM. Red trace is the fitted curve. **(F)** End-point assays of point variants of *Fs*GRE/GRE-AE. 1: WT; 2: C461S; 3: C462S; 4: G785A; 5: C878A; 6: WT + methacrylate; 7: WT + O_2_. Each bar represents mean ± S.D. of triplicate measurements.

We noticed that as-isolated *Fs*GRE/GRE-AE appears as three bands in SDS-PAGE (**Figure 3C**, lane 1). The band between 150 and 100 kDa markers likely corresponds to the full-length protein (126.3 kDa including the His-tag). Because the other two bands appear around 90 and 37 kDa, which add up to the size of full-length protein, they are likely two fragments that arise from protein cleavage. However, anaerobic size-exclusion chromatography (SEC) shows that the protein elutes as one major peak that we assign to a dimeric state (**Figure S5A**), as further supported by a cryo-EM structure described below. This result suggests that the protein is intact in solution under anaerobic conditions and that the fragmentation likely occurs during SDS-PAGE sample preparation. Indeed, the same cleavage is observed in protein gels even for the purest fraction of the major peak from SEC (**Figure S5B**).

We hypothesized that this fragmentation reflects the presence of a glycyl radical cofactor in the as-isolated protein. It is documented that the glycyl radical of GREs reacts with O_2_ to form unstable intermediates, leading to cleavage of the protein backbone (42). If activated *Fs*GRE/GRE-AE undergoes this glycyl radical- and O_2_-mediated backbone cleavage, it would generate protein fragments with molecular weights of 90 and 36 kDa. Since these sizes match the sizes of the fragments that we observe in SDS-PAGE, we hypothesized that we were isolating partially activated *Fs*GRE/GRE-AE from purification under anaerobic conditions and that the protein is cleaved by O_2_ during SDS-PAGE sample preparation. Indeed, the characteristic doublet EPR signal of the glycyl radical can be observed in as-isolated protein sample without *in vitro* activation (**Figure 3D**). Both the EPR signal and the backbone cleavage are absent in the G785A variant that lacks the conserved Gly (**Figure 3C**, lane 4) and in the C878A variant that has inactive GRE-AE domain (**Figure 3C**, lane 5). A similar observation has been made for natively expressed *E. coli* (strain MC4100) PFL, whose O_2_-induced truncation was abolished when the PFL-AE gene was knocked out or the conserved glycine was mutated to an alanine (43). The absence of the glycyl radical in the C878A variant also indicates that the GRE domain cannot be activated by the endogenous PFL-AE in *E. coli* during protein overexpression.

Spin quantification of the glycyl radical signal of as-isolated *Fs*GRE/GRE-AE indicates 0.006 spin per polypeptide, meaning that only 0.6% of the enzyme is activated in the as-isolated state. By screening reductants to enhance the activation yield, Ti^III^-citrate (30, 44) was identified as most reliably producing an increased glycyl radical signal in EPR (5% activation; **Figure S6**) and displaying robust PFL activity (*vide infra*). These results show that *Fs*GRE/GRE-AE contains a functional GRE-AE that can activate the fused GRE partner.

#### *Fs*GRE/GRE-AE shows PFL activity and likely operates through a canonical PFL mechanism

We next assessed whether *Fs*GRE/GRE-AE shows PFL activity. To quantify the activity, as-isolated enzyme or the enzyme activated with Ti^III^-citrate was added into the assay mixture that contains sodium pyruvate and coenzyme A (CoA). Aliquots were taken at multiple timepoints to quantify the acetyl-CoA product using LC–MS/MS. The activated enzyme produces acetyl-CoA with measured kinetic constants of *k*_cat_ = 0.96 ± 0.14 s^−1^ and *K*_M_ = 0.14 ± 0.08 mM for pyruvate (*k*_cat_/*K*_M_ = [6.9 ± 4.0] × 10^3^ M^−1^ s^−1^; **Figure 3E**). The low *K*_M_ value supports that pyruvate is likely a native substrate of the enzyme. Although the observed *k*_cat_ value is orders of magnitude smaller than that of *E. coli* PFL (*k*_cat_ > 100 s^−1^) (45, 46), the catalytic efficiency (*k*_cat_/*K*_M_) of this enzyme is within the range of values observed for previously characterized GREs (47), especially when the value is normalized to the active enzyme concentration (i.e., the glycyl radical concentration) giving the radical-normalized *k*_cat_/*K*_M_ = [1.4 ± 0.8] × 10^5^ M^−1^ s^−1^. 2-ketobutyrate, which is a known substrate of *E. coli* PFL but with lower activity than pyruvate (48), is also metabolized by *Fs*GRE/GRE-AE to produce propionyl-CoA but displays substrate inhibition behavior (**Figure S7**).

To gain insights into the mechanism of the observed PFL activity, we performed activity assays with point variants generated by site-directed mutagenesis (**Figure 3F**). It is known that the glycyl radical is first transferred to the second cysteine (C‘**<u>C</u>**’VS) of the cysteine dyad and then transfers to the other cysteine (‘**<u>C</u>**’CVS). This final thiyl radical then reacts with bound pyruvate to generate a radical adduct that undergoes C–C bond cleavage to liberate formate (**Figure S3A**). We found that both cysteines are indeed required for the activity as both C461S and C462S variants are inactive (**Figure 3F**, samples 2 and 3). Methacrylate, a previously known mechanism-based covalent inhibitor of PFL (49), inhibited wild-type (WT) *Fs*GRE/GRE-AE (**Figure 3F**, sample 6). These observations indicate that *Fs*GRE/GRE-AE performs its PFL reaction via a canonical PFL mechanism involving the conserved cysteine dyad. The G785A variant whose conserved glycine residue is mutated to an alanine and the C878A variant that lacks one of the [Fe_4_S_4_]-binding cysteines are both catalytically inactive (**Figure 3F**, samples 4 and 5). In addition, the WT enzyme is inactivated when the assay is performed aerobically (**Figure 3F**, sample 7). Overall, these results demonstrate the requirement of glycyl radical generation for the PFL activity.

To evaluate the extent to which activation of *Fs*GRE/GRE-AE occurs inter- or intra-molecularly, *Fs*GRE/GRE-AE was activated at varying concentrations (1–10 µM) for activity assays to measure the specific activity: the initial rate of acetyl-CoA production divided by the total enzyme concentration. If the activation occurs primarily through an intramolecular pathway (i.e., the GRE domain is activated by the GRE-AE domain of the same polypeptide), similar amounts of glycyl radical would be installed regardless of the enzyme concentration, leading to similar turnover per polypeptide for samples that are activated at different concentrations. On the other hand, if the activation occurs primarily through an intermolecular pathway (i.e., the GRE domain is activated by the GRE-AE domain of a different molecule), an increase in % GRE activation is expected with increasing *Fs*GRE/GRE-AE concentration and therefore higher turnover per polypeptide would be observed at higher enzyme concentrations. We find that the specific activity of *Fs*GRE/GRE-AE is invariant of the enzyme concentration during activation (**Figure 4**), consistent with intramolecular activation, although we cannot rule out the possibility of intermolecular activation under certain conditions. Collectively, *in vitro* characterization validates *Fs*GRE/GRE-AE as a functional PFL that activates itself intramolecularly, making this fusion protein an ideal system for investigating the structural basis of GRE activation.

**Figure 4.**
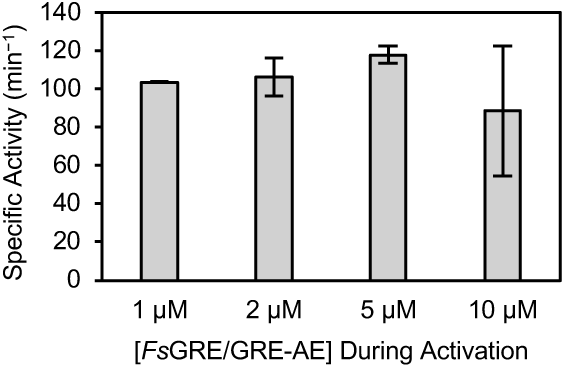
*Fs*GRE/GRE-AE is activated intramolecularly. Measured specific activities of *Fs*GRE/GRE-AE activated at various concentrations. Each bar represents mean ± S.D. of duplicate measurements. Assay conditions: [pyruvate] = 10 mM; [CoA] = 0.1 mM.

### Cryo-EM reveals architecture of a *Fs*GRE/GRE-AE fusion protein

To structurally validate that *Fs*GRE/GRE-AE is a naturally-occurring fusion of a GRE with its cognate AE, we used heterologously expressed *Fs*GRE/GRE-AE to make cryo-EM grids using a blot-free grid preparation method. We collected a dataset containing ∼13,000 movies and processed these movies using cryoSPARC (V4.7.1). Although we were unable to obtain an isotropic map due to preferred orientation of the side view of *Fs*GRE/GRE-AE (**Figure 5A**), our map exhibited clear secondary structural features (**Figure 5B**) which allowed us to fit the *Fs*GRE/GRE-AE AlphaFold 3 (AF3) (50) model as well as the crystal structures of *Ec*PFL (PDB ID 1H18) (51) and *Ec*PFL-AE (PDB ID 3CB8) (7) into the cryo-EM map (**Figure 5C–D**). Our cryo-EM map confirms that this enzyme is a fusion of GRE and GRE-AE domains and supports our SEC data that the oligomeric state of *Fs*GRE/GRE-AE is a homodimer. The dimer interface is between the GRE domains of two *Fs*GRE/GRE-AE protomers, resembling the dimer interface in the *Ec*PFL crystal structure (PDB ID 1H18) (51). On the opposite sides of each GRE from the dimer interface is density for an AE (**Figure 5C–D**). This density is located next to the GRD (**Figure 5E**), exactly where one would expect the AE to bind to install a glycyl radical species in a GRD. Based on the fit of the AF3 model, we appeared to have captured a state of *Fs*GRE/GRE-AE in which the GRD is in its closed position, buried within the GRE active site (**Figure 5C**) and not available for radical installation by the fused AE. This assessment is supported by the fit of *Ec*PFL in our cryo-EM map, that shows density for the GRD in the closed state (**Figure 5D**).

**Figure 5.**
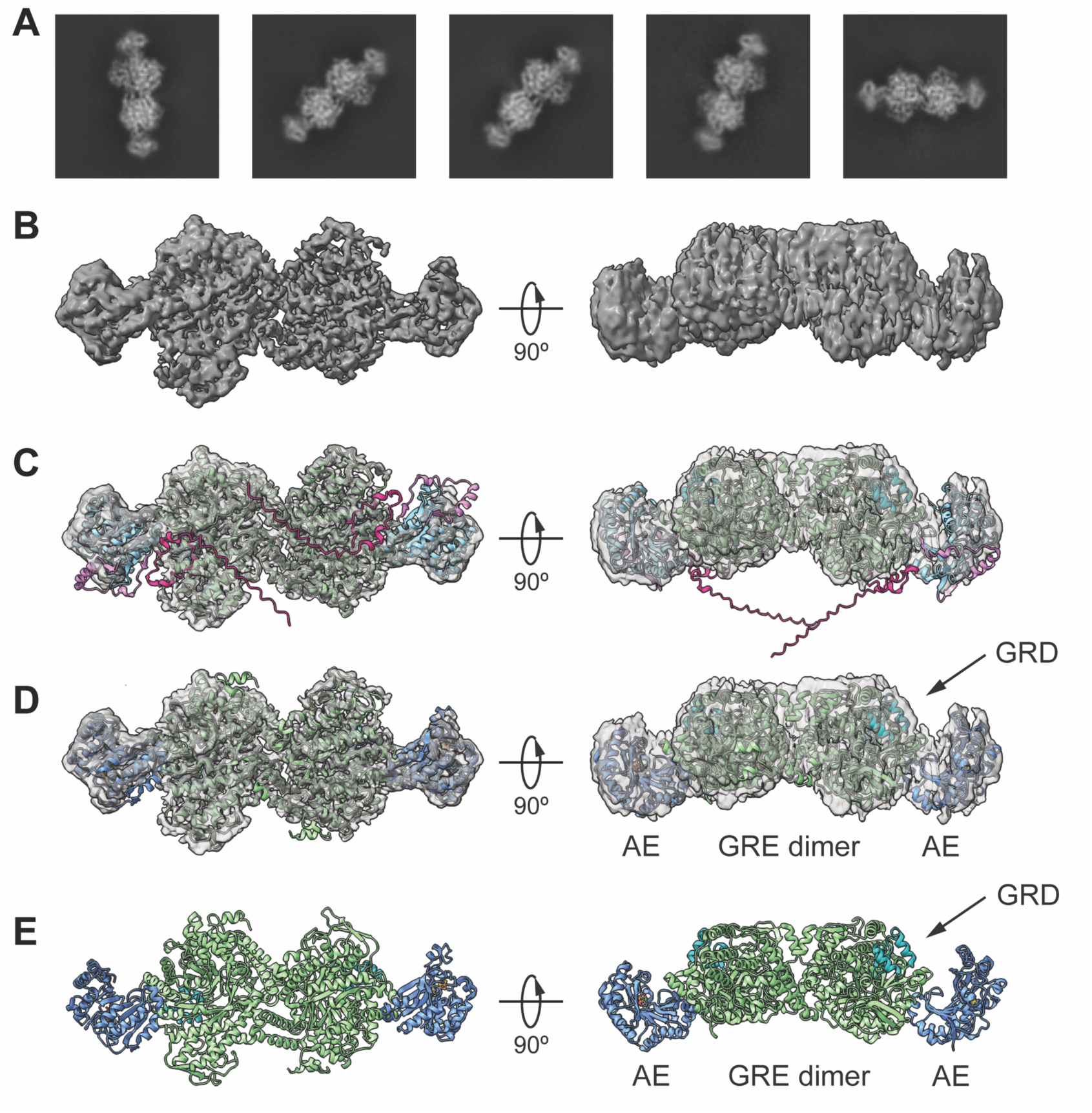
Cryo-EM 2D class averages and 3D reconstruction of *Fs*GRE/GRE-AE reveal the fused architecture. **(A)** Representative 2D class averages show a four-lobed architecture corresponding to a *Fs*GRE/GRE-AE dimer, with the larger lobes corresponding to the GRE domain and the smaller lobes corresponding to the GRE-AE domain. **(B)** The cryo-EM map of dimeric *Fs*GRE/GRE-AE shows some secondary structural features but suffers from preferred orientation as indicated by the streakiness in the 90° rotated view. Isosurface level shown is 0.025. **(C)** An AF3 model of *Fs*GRE/GRE-AE shows adequate fit into the cryo-EM map. The disordered N-terminus (AF3 predicted residues 1-57) is shown in hot pink; GRE domain in green with GRD in teal; linker region in light pink; GRE-AE domain in light blue. **(D)** *Ec*PFL (PDB ID 1H18) and *Ec*PFL-AE (PDB 3CB8) show adequate fit into the cryo-EM map. *Ec*PFL dimer is shown in green with the GRD highlighted in teal. Two copies of *Ec*PFL-AE are shown in blue. **(E)** Structures in (D) with map removed.

### GRE/GRE-AE fusion proteins are widely distributed in diverse microalgae and are expressed in these organisms

To assess whether the observed domain fusions in cluster 45 sequences are a few isolated exceptions or arising from an evolutionarily distinct event, we generated a maximum-likelihood phylogenetic tree of mega-cluster 1 sequences trimmed to the GRE domain (**Figure S8**). GRE/GRE-AE fusion proteins from cluster 45 are found in a single clade within the larger eukaryotic clade. Similar to a previous phylogenetic analysis of PFL sequences (33), eukaryotic PFLs form a single clade adjacent to the *Bacteroidota* clade. These observations indicate that all eukaryotic PFLs share a common ancestor, which is likely derived from an ancestral *Bacteroidota*, and that the fusion of GRE and GRE-AE domains likely occurred as a single evolutionary event in a diatom ancestor.

We then sought to explore the distribution of cluster 45-like GRE/GRE-AE fusion proteins in additional sequence databases. The *Fs*GRE/GRE-AE sequence was used as the query for Basic Local Alignment Search Tool (BLAST) (52) search against the National Center for Biotechnology Information (NCBI) Non-Redundant Protein Database, and hits from this search were used as queries for subsequent rounds of BLAST searching to detect additional homologues. Through this iterative approach, a total of 49 fusion protein sequences were identified predominantly from diatoms but also from closely related algal species, such as alveolate, dinoflagellate, haptophyte, and green algae (**Table S2**). We also note that there are BLAST hits with >60% amino acid sequence identity to GRE/GRE-AE fusion proteins from diatoms but with <70% coverage and annotated as partial sequences; these sequences could also be fusion proteins that currently remain as fragmented sequences.

Diatoms that encode GRE/GRE-AE fusion proteins are taxonomically and ecologically diverse, including both pennate (*Bacillariophycidae*) and centric (*Thalassiosirophyceae*) diatoms and originating from marine, brackish, and freshwater environments around the world (**Figure 6A**). Species in the genera of *Chaetoceros*, *Thalassiosira*, *Skeletonema*, and *Cyclotella* are thought to be globally abundant and widespread in the ocean (53, 54). Organisms such as *Skeletonema costatum* (55), *Mediolabrus comicus* (formerly *Minidiscus comicus*) (56), *Polarella glacialis* (dinoflagellate) (57), and *Prymnesium parvum* (haptophyte) (58) are implicated in algal blooms. Several species including *Fistulifera solaris* (59), *Cyclotella cryptica* (60), and *Skeletonema costatum* (61) are candidate biofuel producers as they are known to accumulate high quantities of fatty acids and triacylglycerol (62). This diversity of encoding organisms indicates potentially wide ecological distribution of GRE/GRE-AE fusion proteins.

**Figure 6.**
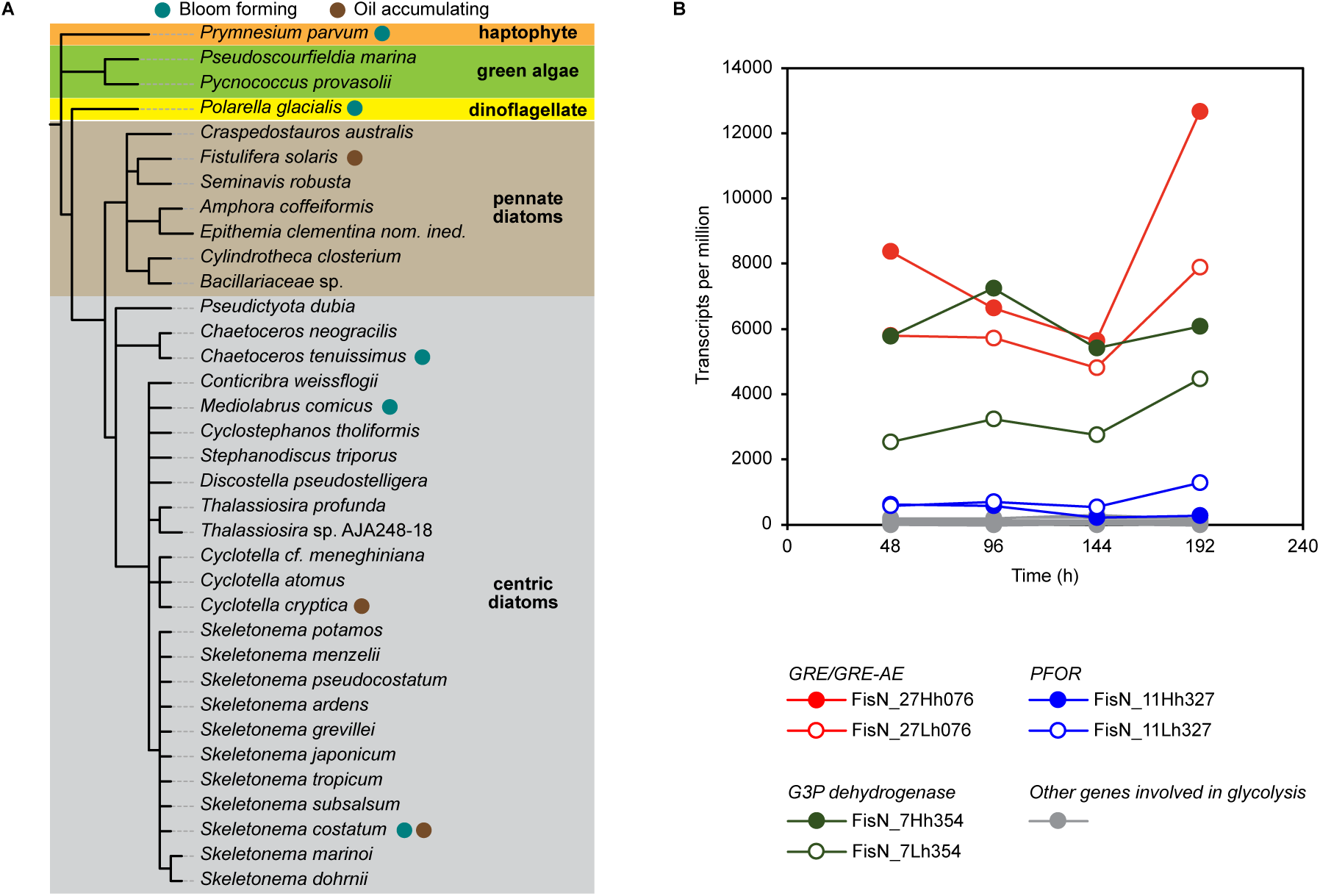
GRE/GRE-AE fusion proteins are widely distributed and expressed in diatoms. **(A)** Taxonomic distribution of organisms that encode GRE/GRE-AE fusion proteins. **(B)** Comparing the expression levels of genes involved in glycolysis and in conversion of pyruvate to acetyl-CoA in *F. solaris* culture (f/2 medium) at different time points (re-analyzed data from ref. 59). PFOR, pyruvate:ferredoxin/flavodoxin oxidoreductase; G3P, glyceraldehyde-3-phosphate.

To evaluate whether genes encoding GRE/GRE-AE fusion proteins are expressed in these organisms, we analyzed publicly available RNA-seq data to quantify transcription levels in transcripts per million (TPM). The TPM values for the fusion protein-encoding genes were compared to those of genes involved in glycolysis, as the fusion proteins are expected to catalyze the conversion of pyruvate to acetyl-CoA. The analysis shows that genes encoding GRE/GRE-AE fusion proteins are expressed at detectable levels in all seven species that we examined (**Figure S9**). In eukaryotes, the conversion of pyruvate to acetyl-CoA is typically mediated by the pyruvate dehydrogenase (PDH) complex, whereas in anaerobic microbes this reaction is commonly catalyzed by a PFL or a pyruvate:ferredoxin/flavodoxin oxidoreductase (PFOR) (63). The encoders of GRE/GRE-AE fusion proteins have genes for PDH and sometimes even for PFOR. For the datasets we analyzed, the expression levels of GRE/GRE-AE fusion proteins and PDH complex subunits were comparable, and in some cases including the oleaginous diatom *F. solaris*, GRE/GRE-AE fusion proteins were more highly expressed than PDH or PFOR (**Figure 6B** and **Figure S9**). Overall, the transcriptomics analysis suggests that GRE/GRE-AE fusion proteins are physiologically relevant and could be key enzymes in pyruvate metabolism by these microalgae.

### Domain architecture analysis reveals additional GRE clusters with potentially novel functions

In addition to cluster 45 GRE/GRE-AE fusion proteins, our SSN and domain architecture analysis identified further uncharacterized clusters that likely possess functions previously unknown in the GRE superfamily. Among the 66 clusters in the AS 200 SSN containing at least 10 unique sequences, seven (cluster 15, 20, 30, 33, 34, 45, and 47) have an average protein length greater than 950 amino acids compared to the average length of 798 amino acids for the whole superfamily (**Table S3**). Cluster 15 contains 86 unique sequences with an average length of 995 amino acids. These sequences have an extension at their N-termini with an SCP2 sterol-binding domain that is implicated in trafficking of lipid molecules (**Figure 7A**) (64). Although most cluster 15 sequences are from uncultured bacteria and archaea, some are from culturable isolates of sulfur-reducing bacteria. Cluster 15 sequences belong to mega-cluster 2 in the AS 100 SSN and are part of the ‘dehydratase-type’ cluster in the AS 150 SSN (**Figure S10A**). Genomic neighborhoods of these sequences contain genes encoding a highly conserved putative GRE-AE and a moderately conserved short chain dehydrogenase annotated as 3-dehydrosphinganine reductase. Together, these observations suggest that these GRE/SCP2 fusion proteins could be involved in a C–O bond cleavage reaction of a lipid substrate, an activity currently unknown for GREs.

**Figure 7.**
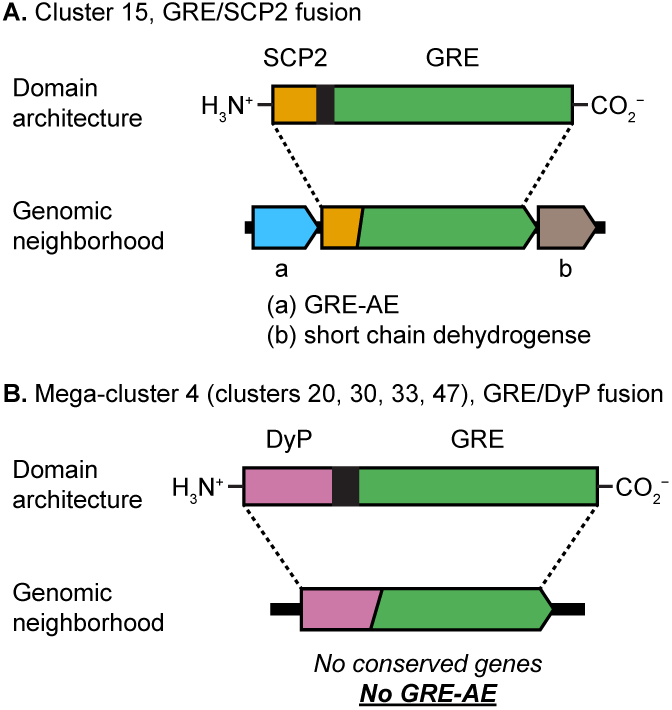
Domain analysis reveals additional GRE fusion proteins. **(A)** Cluster 15 contains GRE/SCP2 fusion proteins with a highly conserved GRE-AE and a moderately conserved short chain dehydrogenase annotated as “3-dehydrosphinganine reductase” in their genomic neighborhoods. **(B)** Mega-cluster 4 contains GRE/DyP fusion proteins with no GRE-AE in their genomic neighborhoods.

Cluster 34 sequences have an average length of 987 amino acids, but no additional domain can be assigned. The remaining four clusters (cluster 20, 30, 33, and 47) contain sequences with average lengths of 1100–1300 amino acids and they all belong to mega-cluster 4 in the AS 100 SSN (**Figure S10B**). Most sequences of mega-cluster 4 contain a dye-decolorizing peroxidase (DyP) domain fused to the GRE domain (**Figure 7B**), while some are standalone GREs with a DyP-encoding gene as the genomic neighbor. Unexpectedly, genes that encode GRE-AEs are absent in genomic neighborhoods of mega-cluster 4 sequences. Although a few of these GRE/DyP fusion proteins have been previously noted (65–67), their broader distribution in the GRE superfamily has been unknown and they remain uncharacterized. These GRE/DyP fusion proteins appear in a wide range of aerobic organisms from bacteria, fungi, and metazoa, including pathogens of humans, plants, and fungi such as *Burkholderia cepacia*, *Fusarium proliferatum*, *Ralstonia solanacearum*, and *Janthinobacterium agaricidamnosum*. Because these fusion proteins belong to the uncharacterized mega-cluster 4, have no adjacent GRE-AE, and are found in primarily aerobic species, we hypothesize that they have a unique activity that enables GRE-mediated chemistry in more oxidizing environments. The identification of these novel GRE fusion proteins further demonstrates domain architecture analysis as a productive strategy for enzyme discovery.

## Discussion

GREs have diverse functions that impact primary metabolism of obligate and facultative anaerobes. Although there are ∼20 unique reactions of GREs that have been characterized, this number likely represents only a small fraction of the full functional repertoire of the superfamily given its high sequence diversity. While recent advances in DNA sequencing technology have significantly expanded the number of putative enzymes, navigating the vast sequence space and prioritizing genes for biochemical characterization are continued challenges in the field (16, 23–25). Through our SSN analysis of the GRE superfamily (PF02901), we have surveyed the sequence and functional diversity and rapidly identified clusters of unknown function. Of the 66 clusters with ≥10 unique sequences, only 15 clusters had at least one biochemically characterized enzyme. By varying the alignment score threshold of SSN, sequence similarity and hence functional relationships of these GRE clusters are revealed. Based on how they form mega-clusters with characterized GREs at low AS threshold, the general activity of many uncharacterized clusters can be predicted. Clusters that instead form distinct mega-clusters likely represent reaction classes not yet known in the GRE superfamily. This analysis serves as the first step in predicting the reactivity of unknown clusters for functional characterization. We expect that this type of analysis will benefit from periodic revisiting as sequence databases continue to grow and as more enzymes are biochemically characterized.

In this study, we analyzed domain architecture to identify clusters containing unusual domain fusions. This strategy has been used previously to find novel enzymes in the crotonase superfamily (22), the monotopic phosphoglycosyl transferase superfamily (31), and the 4-oxalocrotonate tautomerase superfamily (32), as well as unusual polysaccharide monooxygenases (68) and bifunctional terpene synthases (69). Using this approach, we identified three previously unappreciated domain fusions of GREs. Cluster 45 contains GRE/GRE-AE fusion proteins from diverse diatoms and related organisms and is part of the PFL mega-cluster with cluster 1 (canonical PFLs) and cluster 5 (*Streptococcaceae* PFLs). Our biochemical characterization of a GRE/GRE-AE fusion from the diatom *F. solaris* has shown that the enzyme is a functionally active PFL that appears to operate via canonical PFL mechanism, consistent with the prediction from our SSN analysis. The physiological role of these GRE/GRE-AE fusion proteins in diatoms and the functional relevance of the domain fusion are currently under investigation. We speculate that the domain fusion could contribute to enhanced effective concentrations of the GRE and GRE-AE domains and efficient co-regulation of transcription and translation of the two genes (70).

A unifying mechanistic feature of all GREs characterized to date is that an oxygen-sensitive glycyl radical cofactor must be installed post-translationally by a separate GRE-AE that is specific to each GRE. The glycyl radical installation by a GRE-AE is thought to require a large conformational change of the cognate GRE. Due to the lack of structural information on the GRE activation process, how a GRE and its GRE-AE interact specifically with each other to induce such a large structural motion remains elusive. We believe the GRE/GRE-AE fusion protein identified and characterized herein serves as an excellent model for studying this interaction. Our current low resolution cryo-EM map shows the overall dimeric architecture of *Fs*GRE/GRE-AE fusion protein. The crystal structures of *Ec*PFL and *Ec*PFL-AE fit adequately into the map, with density surrounding the GRD (**Figure 5D**). The observed orientations of *Ec*PFL and *Ec*PFL-AE (**Figure 5E**) support the previous hypothesis that the GRD swings out of the buried PFL active site to become accessible to PFL-AE (**Figure S11**) (7).

The identification of GRE/GRE-AE fusion protein genes from taxonomically diverse diatoms and the high transcriptional abundance of these genes from several diatom species indicate that these enzymes could play an important role in pyruvate metabolism in these organisms. This finding is notable because there are limited examples of GREs in eukaryotes and most of them are from anaerobic protists (35–37). There is one example of PFL in the green alga *C. reinhardtii* that is expressed inside the mitochondrion or chloroplast under anoxic conditions (38, 39). While diatoms are generally considered as aerobes, they have broad metabolic capabilities that enable their survival under anoxic conditions (34, 63, 71). Our identification and characterization of cluster 45 further expands the metabolic versatility of diatoms and the physiological role of GREs beyond anaerobic bacteria.

In conclusion, we have leveraged SSN analysis to survey the functional diversity of the GRE superfamily and identified a unique cluster of GRE/GRE-AE fusion proteins from diverse photosynthetic microalgae that are functional PFLs. Using this native fusion, we have obtained preliminary structural information on the interaction between GRE and GRE-AE by cryo-EM, but an isotropic high-resolution structure will be needed to better understand the molecular basis of glycyl radical installation. Functional characterization of the two additional GRE fusion proteins identified in this study, GRE/SCP2 fusion proteins from cluster 15 and GRE/DyP fusion proteins from mega-cluster 4, is currently underway. This work demonstrates the utility of SSN and domain architecture analysis in identifying novel enzymes from enzyme families with high sequence diversity, and we anticipate that the workflow employed in this study will be applicable to other large, functionally diverse enzyme families.

## Materials and Methods

### General considerations

All reagents were purchased from commercial suppliers and used without further purification unless otherwise noted. The plasmids constructed in this work were deposited into Addgene. Computational analyses were performed on the Cannon Cluster of the Harvard Faculty of Arts and Sciences Research Computing. Experimental details on phylogenetic tree construction, transcriptomic analysis, SEC, EPR spectral analysis, and LC–MS/MS are described in the Supporting Information (SI).

### SSN construction

SSNs were generated using EFI-EST,(24, 25) Option B with PF02901 (UniProt version 2025-03, InterPro version 106) as the input family, excluding fragment sequences. All-by-all BLAST was performed with sequences trimmed to PF02901 domain. Networks were subsequently generated with alignment score thresholds of 100, 110, 115, 120, 125, 150, and 200. The resulting networks were visualized with Cytoscape 3.10 (72).

### Plasmid construction

pET28a–*Fs*GRE/GRE-AE, the plasmid encoding full-length *Fs*GRE/GRE-AE (UniProt ID: A0A1Z5KPM5) between *Nde*I and *Xho*I restriction sites of pET28a(+) vector, was ordered from GenScript. Site-directed mutagenesis was performed by first amplifying N- and C-terminal fragments through PCR using Q5 High-Fidelity Master Mix (New England Biolabs) with primers ordered from GENEWIZ (**Table S4**) and then ligating the fragments to restriction-digested pET28a(+) backbone through Gibson assembly using NEBuilder HiFi DNA Assembly Master Mix (New England Biolabs). Sequences of the resulting constructs (pET28a–*Fs*GRE/GRE-AE–C461S, –C462S, – G785A, and –C878A) were validated through whole plasmid sequencing by Plasmidsaurus.

### Heterologous expression

Chemically competent *E. coli* BL21(DE3) cells were transformed with the constructed plasmid using a standard protocol. The transformed cells were streaked on an LB/kanamycin plate, and the plate was incubated at 37 °C overnight. A 200 mL LB starter culture with 50 µg/mL kanamycin was inoculated with a single colony from the plate and shaken at 37 °C, 200 rpm for 18 h. 2 L of LB medium containing 50 µg/mL kanamycin was inoculated with 20 mL of starter culture. The culture was supplemented with 25 µM FeCl_3_ and shaken at 37 °C, 180 rpm. At OD_600_ = 0.3, the culture was supplemented with 300 µM L-cysteine and an additional 25 µM FeCl_3_. At OD_600_ = 0.6, the culture was cooled in an ice bath before the expression was induced with 500 µM IPTG. The culture was shaken at 18 °C, 180 rpm for 18 h. Cells were harvested by centrifugation at 8,000 × *g*, 4 °C for 5 min. The resulting cell pellet was flash frozen in liquid N_2_ and stored at −80 °C until further use.

### Protein purification

All purification steps were performed in an anaerobic chamber (Coy Labs) containing 97% N_2_ and 3% H_2_ (<5 ppm O_2_) housed inside a 4 °C cold room. Cells were resuspended in 3 mL per g cell paste of 25 mM HEPES, pH 7.5, 10% v/v glycerol, 200 mM NaCl, 2 mM NaDT, 1 mg/mL lysozyme, 0.1 mg/mL DNase I, and EDTA-free Pierce protease inhibitor tablets. After the resuspension, cells were lysed by sonication at 10% amplitude for 5 min (10 s on followed by 25 s off). The lysate was clarified by centrifugation at 20,000 × *g*, 4 °C for 1 h. The cleared lysate was loaded onto a Ni-NTA column, equilibrated with 25 mM HEPES, pH 7.5, 10% v/v glycerol, 200 mM NaCl, 2 mM NaDT. The resulting column was washed with 5 column volumes (CVs) of 25 mM HEPES, pH 7.5, 10% v/v glycerol, 200 mM NaCl, 2 mM NaDT, 20 mM imidazole. The protein was eluted with at least 3 CVs of 25 mM HEPES, pH 7.5, 10% v/v glycerol, 200 mM NaCl, 2 mM NaDT, 300 mM imidazole. The fractions containing the protein, determined by dark brown color or SDS-PAGE, were combined and concentrated to 2 mL using Ultra-15 Centrifugal Filters with a 50 kDa MWCO. The concentrated protein was oxidized with 0.5 mL of 10 mM indigodisulfonic acid (IDS) to poise the [Fe_4_S_4_] cluster at 2+ oxidation state, and the excess IDS was removed by centrifugation at 7,000 × *g* for 10 min and gel-filtration through a PD-10 column (Cytiva) equilibrated with 25 mM HEPES, pH 7.5, 10% v/v glycerol, 200 mM NaCl. Purity of the protein was determined by SDS-PAGE. Concentration of the protein was estimated using the Pierce BCA assay kit (Thermo Fisher Scientific). Fe content of the protein was determined by a ferene assay (73). The protein was aliquoted, frozen in liquid N_2_, and stored in liquid N_2_ until further use.

### Activation of FsGRE/GRE-AE

All steps were performed in an MBraun UNILAB glovebox under a N_2_ atmosphere (<5 ppm O_2_). To a solution of *Fs*GRE/GRE-AE (1–10 µM for activity assay, 100 µM for EPR spectroscopy) in standard buffer (25 mM HEPES, pH 7.5, 10% v/v glycerol, 200 mM NaCl) were added AdoMet (as 10 mM stock in standard buffer) and a reductant (NaDT or Ti^III^-citrate; as 100 mM stock in standard buffer; 100 mM Ti^III^-citrate stock solution was prepared by mixing equal volume of 200 mM TiCl_3_·(thf)_3_ and 200 mM sodium citrate tribasic salt), both to a final concentration of 1 mM. The resulting mixture was incubated at room temperature for 30 min before it was used for EPR spectroscopy or activity assays.

### EPR spectroscopy

200 µL of as-isolated or activated *Fs*GRE/GRE-AE was transferred to a clear-fused quartz EPR tube (Wilmad) inside an MBraun UNILAB glovebox, capped with a rubber septum, and frozen in liquid N_2_ outside the glovebox. EPR spectra were recorded on a Bruker ElexSys E500 spectrometer at the Laukien-Purcell Instrumentation Center (Department of Chemistry and Chemical Biology, Harvard University). 1 mM CuSO_4_ with 10 mM HCl and 2 M NaClO_4_ was used as the standard for spin quantification (74).

### Activity assay

The procedure was adapted from previous studies (46, 75). Assays were set up in an MBraun UNILAB glovebox under a N_2_ atmosphere (<5 ppm O_2_). Assays were performed in duplicates or triplicates. 20 µL of as-isolated *Fs*GRE/GRE-AE or 20 µL of activated *Fs*GRE/GRE-AE (1–10 µM monomer) that has been diluted 10-fold with standard buffer was added to a 180 µL solution of assay mixture containing 10 mM sodium pyruvate (Sigma-Aldrich) and 0.1 mM coenzyme A trilithium salt (RPI) in standard buffer. The mixture was incubated at room temperature for certain periods of time before a 20 µL aliquot was taken out from the glovebox and quenched with 2 µL of LC–MS grade acetic acid (Millipore Sigma). The quenched aliquots were stored at −80 °C until further use. Acetyl-CoA was quantified using LC–MS/MS (see SI for details).

### Michaelis–Menten kinetics

10 µM *Fs*GRE/GRE-AE was activated with Ti^III^-citrate following the procedure described above. Analogous to the activity assay procedure described above, the activated enzyme was first diluted 10-fold with standard buffer before 20 µL of the diluted mixture was added to 180 µL solutions of assay mixtures containing 0.05, 0.1, 0.15, 0.2, 0.3, 0.4, 0.6, 0.8, 1, 2, 5, or 10 mM sodium pyruvate and 0.1 mM CoA. Aliquots were taken for quantification of acetyl-CoA using LC–MS/MS.

### Cryo-EM sample preparation, data collection, and processing

A frozen *Fs*GRE/GRE-AE (containing an N-terminal His_6_-tag) aliquot in standard buffer and a 100 mM AdoMet (Sigma Aldrich) stock (in 20 mM HEPES, pH 8) was brought into an MBraun anaerobic chamber at 4 °C under a N_2_ atmosphere with <1 ppm O_2_. The protein was buffer exchanged into anoxic grid buffer (25 mM HEPES (Sigma-Aldrich), pH 7.5, 200 mM NaCl (Fisher Chemical)). Concentration was assessed by measuring absorbance at 280 nm using a NanoDrop 2000c spectrophotometer (Thermo Scientific) with the following parameters: molecular weight = 126.32 kDa and ε/1000 = 128.58 M^-1^ cm^-1^. In an Eppendorf tube, *Fs*GRE/GRE-AE was diluted to 1 mg/mL in grid buffer, and the 100 mM AdoMet stock was added to a final concentration of 200 µM. This sample was sealed inside of a glass bottle until grid plunging. An SPT Labtech chameleon automatic cryo-plunging system was used to prepare a holey carbon-coated copper Quantifoil Active™ cryo-EM grid (300 mesh, hole size 1.2 µm, hole spacing 0.8 µm; batch ID: CG001587). The protocol for chameleon grid preparation had previously been optimized for oxygen-sensitive samples (76, 77). The grid was glow discharged at −12 mA for 60 s and plunged into liquid nitrogen-cooled ethane (−174 °C) with a final dispense-to-plunge time of 151 ms. The plunge temperature was 24.3 °C and plunge relative humidity was 78%. The grid was screened on a Talos Arctica microscope at the Cryo-EM Facility at MIT.nano.

Data collection information is summarized in the data collection table (**Table S5**). Data were collected at Pacific Northwest Cryo-EM Center (PNCC) using a Thermo Fisher Titan Krios 300 keV transmission electron microscope equipped with a Falcon 4i direct electron detector and Selectris X imaging filter at 215,000 x magnification (0.5727 Å/pixel). A total of 13,304 movies were collected with a defocus range of −0.5 to −2.0 μm and a total exposure of 50 e^−^/Å^2^.

The cryo-EM data processing was performed using cryoSPARC V4.7.1 (78) (**Figure S12**). The movies were imported using 40 EER fractions and an EER upsampling factor of 1 (no upsampling). Movies were motion-corrected using patch motion correction followed by patch contrast transfer function (CTF) estimation. The pretrained ResNet16 Topaz model picked on all micrographs (79). The ∼573,000 picked particles were extracted with a box size of 576 and downsampled to a box size of 192 (1.718 Å/pixel). The particles were 2D classified (200 classes) and a set of three of the resulting class averages displaying “side views” (∼28,000 particles boxed in magenta) and another set (∼9,000 particles boxed in teal) displaying possible “top views” were selected to train two separate Topaz models for autopicking. The resulting Topaz models were used to pick particles from the entire set of micrographs, and each set of Topaz picked particles (∼460,000 top view or 1.3 million side view particles) were extracted with a box size of 576 and downsampled to a box size of 192 (1.718 Å/pixel). Particles from each set were 2D classified (200 classes). The chosen classes from each magenta (side view) and teal (top view) set were combined to form a particle stack of ∼1.2 million particles, which were chosen for *ab initio* reconstruction (4 classes). Two classes (total of ∼764,000 particles) were chosen for 3D classification (10 classes) with a filter resolution of 40 Å. Eight classes (∼632,000 particles) were chosen for one *ab initio* reconstruction (only ∼218,000 particles used) and was subsequently refined with non-uniform refinement (NUR) with a reported resolution of 3.53 Å (∼632,000 particles). These particles were re-extracted using box size of 576 to retain the data collection pixel size (0.5727 Å/pixel). (Some particles could not be re-extracted with new box size because they were too close to the edge of the micrograph.) The re-extracted particles (∼619,000) were used for a NUR of the *ab initio* reconstruction which resulted in a 3.18 Å-resolution reconstruction. These particles were then 3D classified (10 classes) with a filter resolution of 60 Å. Seven classes were chosen to perform NUR with minimization over per-particle scale on the previous 3.18 Å-resolution reconstruction, which resulted in a new 3.2 Å-resolution reconstruction (∼423,000 particles) and a conical Fourier shell correlation area ratio (cFAR) score of 0.02, as calculated by cryoSPARC, which indicates preferred orientation (**Figure S13A–B**). The map’s corresponding viewing direction distribution plot from cryoSPARC also indicates that a major portion of the dataset only represents a narrow set of views, as shown by the isolated red hotspots (**Figure S13C**). For another assessment of preferred orientation, the final cryo-EM map, mask, and half maps were used for 3D FSC calculation using the Salk Institute for Biological Sciences remote 3D FSC server (https://3dfsc.salk.edu/), which reports the map at 3.27 Å global resolution and 0.746/1 sphericity score **(Figure S13D**). The directional FSC curves indicate that the map suffers from severe anisotropy, with the x direction being poorly resolved. An AlphaFold 3 model of *Fs*GRE/GRE-AE and crystal structures of PFL (PDB ID 1H18) (51) and PFL-AE (PDB ID 3CB8) (7) were fit into the map using UCSF ChimeraX version 1.11.1 (80).

## Acknowledgements

This work was completed in part with resources at the Laukien-Purcell Instrumentation Center of the Department of Chemistry and Chemical Biology at Harvard University. The computations in this paper were run on the FASRC Cannon cluster supported by the Faculty of Arts and Sciences Division of the Science Research Computing Group at Harvard University. We thank Dr. Grace Kenney for initial assistance with the SSN analysis and Dr. Ruocheng Yu for helpful discussions. Cryo-EM grids were prepared in the Department of Biology Cryo-EM Facility in MIT.nano using a SPT Labtech chameleon, which was purchased using funds from the HHMI Transformative Technology 2019 Award. Cryo-EM grids were screened on the Talos Arctica microscope in the Department of Biology Cryo-EM Facility in MIT.nano, which was a gift from the Arnold and Mabel Beckman Foundation. A portion of this research was supported by the National Institutes of Health grant R24GM154185 and performed at the Pacific Northwest Center for Cryo-EM (PNCC) with assistance from Dr. Omar Davulcu. This work was additionally supported by the National Institutes of Health grant R35GM126982 (to CLD). CLD and EPB are Howard Hughes Medical Institute Investigators. The content is solely the responsibility of the authors and does not necessarily represent the official views of the National Institutes of Health.

## Author Contributions

G.N. and E.P.B. designed research; G.N. and N.M.G. performed research; G.N., N.M.G., C.L.D., and E.P.B. analyzed data; G.N., N.M.G., C.L.D., and E.P.B. wrote the paper.

