## Supporting Information for "Domain Architecture Analysis Guides the Discovery of Unusual Glycyl Radical Enzyme Fusion Proteins"

**Table of Contents:**

A. Supplementary Methods ................................................ S2

B. Supplementary Figures ................................................ S4

C. Supplementary Tables ................................................ S17

D. Supplementary References ................................................ S22

### A. Supplementary Methods

*Construction of a maximum-likelihood phylogenetic tree of mega-cluster 1 sequences*. To generate the multiple sequence alignment (MSA) of mega-cluster 1 sequences, fragments were first removed using Seqtk[1] by filtering out sequences shorter than 200 amino acids before the sequences were clustered to 80% amino acid identity using CD-HIT[2, 3] and aligned using MAFFT[4, 5] with the automatic model. The resulting MSA was trimmed to remove ambiguities using the trimAl automated algorithm.[6] A maximum-likelihood phylogenetic tree was constructed from the trimmed MSA using IQ-TREE2[7] with automated ModelFinder[8] and 1000 ultrafast bootstraps.[8] The resulting tree was visualized using ggtree.[9]

*Transcriptomic analysis*. The following RNA-seq datasets were downloaded from the European Nucleotide Archive (ENA): PRJDB2969, *Fistulifera solaris*;[10] PRJEB36275, *Seminavis robusta*;[11] PRJNA217458, *Cylindrotheca closterium*; PRJNA310432, *Skeletonema marinoi*; PRJEB76158, *Prymnesium parvum*; PRJNA589195, *Cyclotella cryptica*;[12] PRJDB10789, *Chaetoceros tenuissimus*. The following transcriptomes were downloaded from the annotated genomes of each organism from the NCBI Genome Database: GCA_002217885.1, *Fistulifera solaris*; GCA_903772945.1, *Seminavis robusta*; GCA_933822405.4, *Cylindrotheca closterium*; GCA_030871285.1, *Skeletonema marinoi*; GCA_041296205.1, *Prymnesium parvum*; GCA_013187285.2, *Cyclotella cryptica*; GCA_021927905.1, *Chaetoceros tenuissimus*. Salmon[13] (version 1.10.2) was used to index the transcriptome and quantify the RNA-seq reads for each gene, resulting in estimated transcript abundance in transcripts per million (TPM).

*Size-exclusion chromatography (SEC)*. SEC was performed inside an anaerobic chamber (Coy Labs) using an AKTA Pure fast protein liquid chromatography system (Cytiva) equipped with a UV detector (280 nm) and a Superdex 200 HiLoad 16/60 column (Cytiva). The column was washed with 2 column volumes (CVs) of water containing 2 mM sodium dithionite (NaDT) and equilibrated with 2 CVs of running buffer (25 mM HEPES, pH 7.5, 10% v/v glycerol, 200 mM NaCl) prior to sample injection (250 µL of 25 µM *Fs*GRE/GRE-AE). The protein was eluted with 1.3 CVs of running buffer at a flow rate of 0.5 mL/min. 5 mL fractions were collected. Gel Filtration Standard #1511901 (Bio-Rad) was used to estimate the size of the protein.

*EPR spectral simulation and spin quantification*. EPR spectra were simulated using EasySpin[14] (version 6.0.7) *pepper* function in MATLAB (version R2024b). Spin concentration was estimated based on a previously reported method[15] using 1 mM $CuSO_4$ with 10 mM HCl and 2 M $NaClO_4$ as the standard. Baseline correction and double integration of experimental spectra were performed in MATLAB (version R2024b).

*End-point activity assays of FsGRE/GRE-AE variants or in the presence of inhibitors.* WT, C461S, C462S, G785A, or C878A variants were activated and added into activity assay mixture following the procedure described in the main text. For methacrylate inhibition: 10 µM activated WT enzyme was first diluted 10-fold into standard buffer containing 10 mM methacrylic acid. The resulting mixture was incubated for 10 min before it was added into the assay mixture. For $O_2$ inhibition: 10 µM activated WT enzyme was taken out from the glovebox and diluted 10-fold into aerobic standard buffer. The diluted mixture was incubated for 30 min before it was added into the assay mixture. The assay mixtures were incubated for 2 h before aliquots were taken for acetyl-CoA quantification using LC–MS/MS.

*Enzyme kinetics with 2-ketobutyrate.* 20 µL of 10 µM as-isolated *Fs*GRE/GRE-AE was added into 180 µL of assay mixture containing 0.1 mM CoA and varying concentrations of 2-ketobutyrate (0.05, 0.1, 0.2, 0.3, 0.4, 0.6, 0.8, 1, 2, 3, 5, and 10 mM) in standard buffer. 20 µL aliquots were taken at each timepoint for quantification of propionyl-CoA using LC–MS/MS.

*Acetyl-CoA quantification using LC–MS/MS*. The procedure was adapted from a previous study.[16] Quenched aliquots from activity assays were thawed at room temperature and diluted 10-fold with LC–MS grade water containing 5 µM $d_3$-acetyl-CoA sodium salt (Cayman) as an internal standard. The diluted mixtures were incubated at −20 °C for 15 min before they were filtered through a 0.2 µm wwPTFE filter plate (Pall Corp.). The filtrates were further diluted 5-fold with LC–MS grade water and kept at 4 °C prior to LC–MS/MS analysis. Standards were prepared according to the same methods used for experimental samples. LC–MS/MS analysis was performed using a Waters Acquity UPLC H-Class System coupled to Water Xevo TQ-S (Waters Corp., Milford, MA). Liquid chromatography was conducted using the following method: Solvent A = 30 mM ammonium acetate in water. Solvent B = 0.1% v/v formic acid in acetonitrile. The samples (2 µL) were injected onto a Waters Acquity UPLC BEH C18 column (2.1 × 50 mm, 1.7 µm) equilibrated with 95% Solvent A, flow rate of 0.8 mL/min at 40 °C. The following gradient was applied: (i) 0–0.5 min at 100% A isocratic, (ii) 0.5–4.0 min at 0–20% B gradient, (iii) 4.0–4.1 min at 20–100% B gradient, (iv) 4.1–4.4 min at 100% B isocratic, (v) 4.4–4.5 at 100–0% B gradient, and (vi) 4.5–5.0 min at 100% A isocratic. MS/MS detection was performed using electrospray ionization in positive mode (ESI+) with the following source parameters: capillary voltage, 0.50 kV; cone voltage, 85 V; source offset voltage, 50 V; desolvation temperature, 600 °C; desolvation gas flow, 1000 L/h; cone gas flow, 150 L/h; nebulizer, 7.0 bar. The following MS/MS transitions and collision energies were used to detect acetyl-CoA and $d_3$-acetyl-CoA: (i) acetyl-CoA: parent ion $m/z$ = 810.41; daughter ion $m/z$ = 303.22; collision energy = 30 V; cone voltage = 66 V. (ii) $d_3$-acetyl-CoA: parent ion $m/z$ = 813.43; daughter ion $m/z$ = 306.26; collision energy = 34 V; cone voltage = 66 V.

*Propionyl-CoA quantification using LC–MS/MS*. The same procedures for acetyl-CoA quantification were used, except that propionyl-CoA lithium salt (Chem-Impex) was used as the standard (with $d_3$-acetyl-CoA as an internal standard) and that the following MS/MS transitions and collision energies were used: (i) propionyl-CoA: parent ion $m/z$ = 824.49; daughter ion $m/z$ = 317.24; collision energy = 32 V; cone voltage = 72 V. (ii) $d_3$-acetyl-CoA: parent ion $m/z$ = 813.44; daughter ion $m/z$ = 306.25; collision energy = 32 V; cone voltage = 16 V.

### B. Supplementary Figures

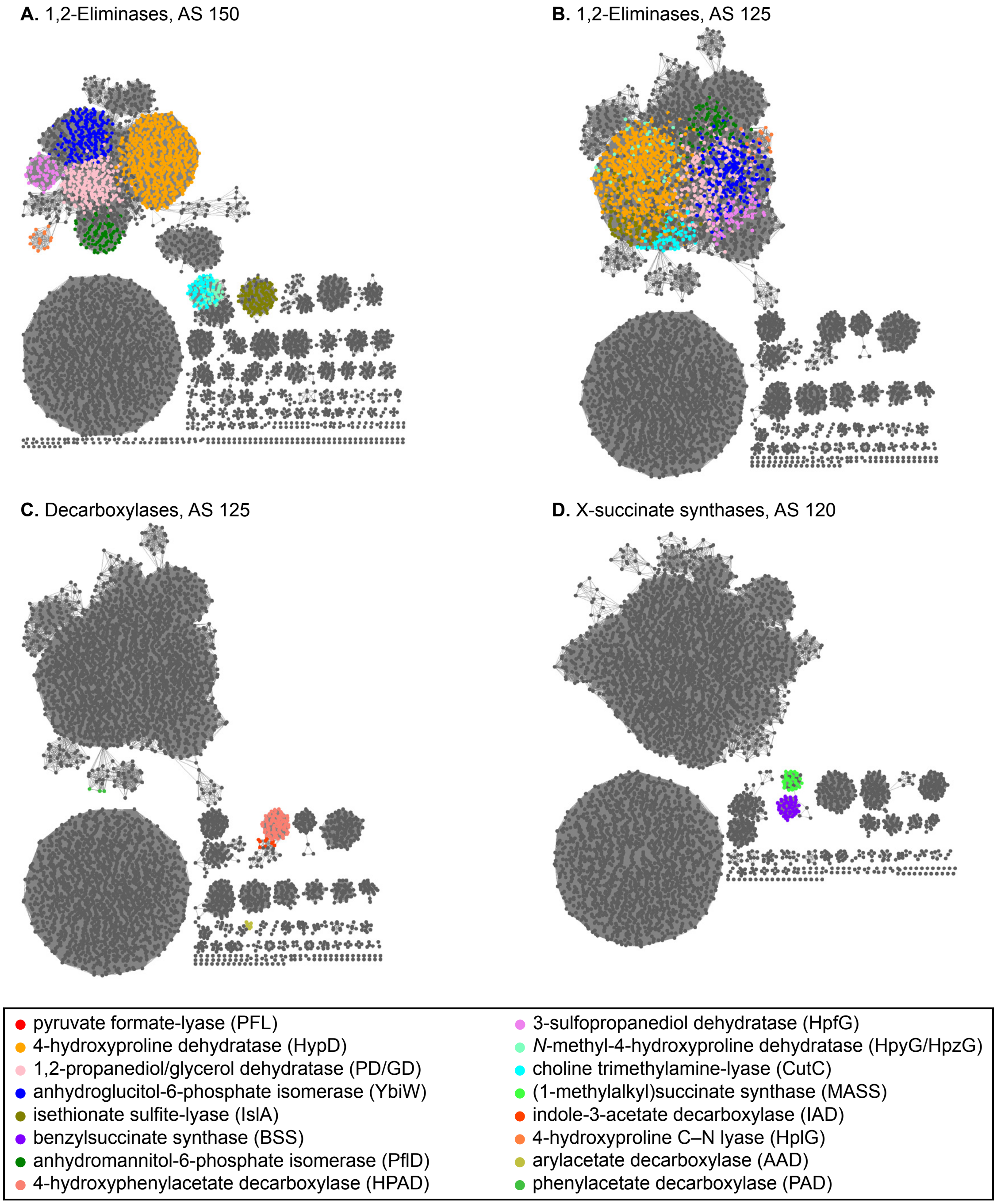


**Figure S1.** SSNs of PF02901-containing sequences at varying alignment score (AS) thresholds. Isofunctional clusters from the AS200 SSN are colored as described in figure. **(A, B)** At AS threshold 150, the clusters corresponding to YbiW, HypD, PflD, PD, GD, and HpfG merge together, and at AS threshold of 125, CutC, HpyG, HpzG, and IslA clusters are also merged to form the larger 1,2-eliminase cluster. **(C)** At AS threshold of 125, HPAD and IAD clusters form the larger decarboxylase cluster. **(D)** At AS threshold of 120, BSS and MASS clusters form the larger XSS cluster.

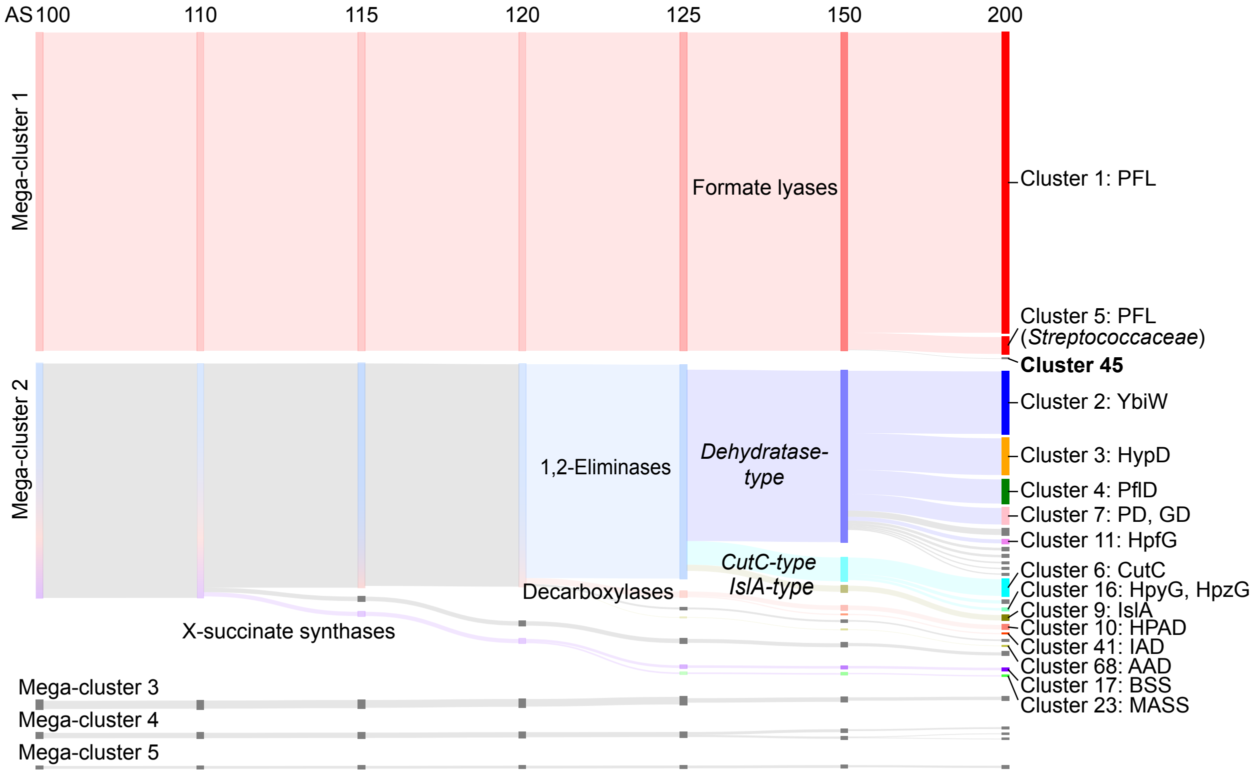


**Figure S2.** A Sankey diagram showing the relationship of GRE clusters as a function of the AS threshold. This diagram quickly reveals the GRE class (formate lyase, 1,2-eliminase, decarboxylase, X-succinate synthase) of a given uncharacterized cluster.

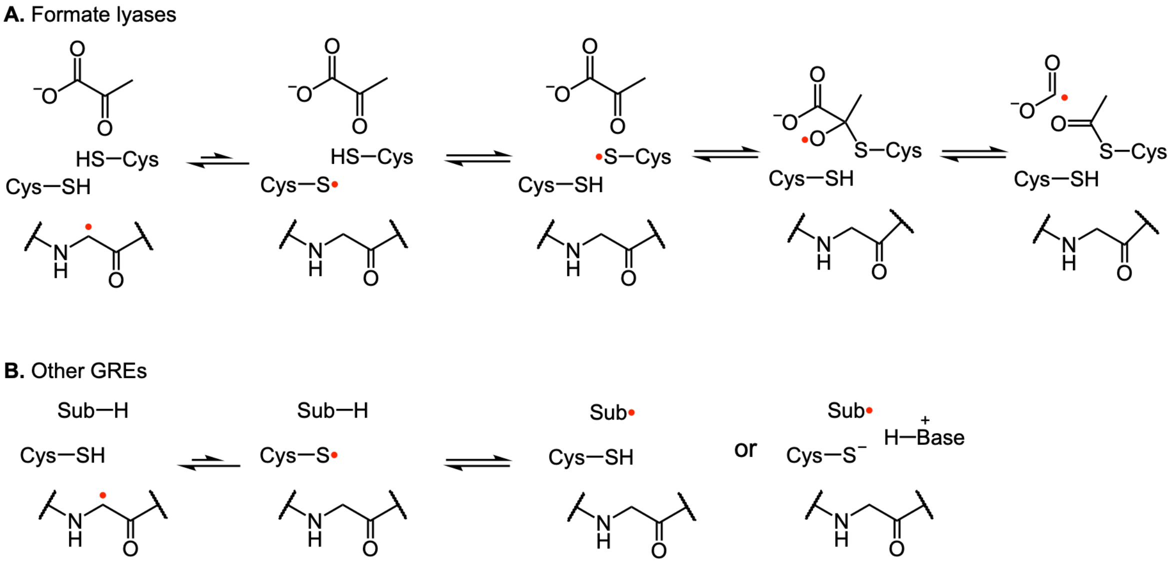


**Figure S3.** Thiyl radical reactivities of GREs. **(A)** Formate lyases. The glycyl radical is first transferred to the conserved cysteine residue and then transferred to the second cysteine conserved within this class. The second cysteine is the cysteine that reacts with substrate (in this case, pyruvate), and the reaction involves formation of a covalent adduct. **(B)** Other GREs. The glycyl radical is transferred to the conserved cysteine residue. The thiyl radical abstracts a H-atom from the substrate (H-atom transfer) or accepts an electron from the substrate, often concurrent with a proton transfer (proton-coupled electron transfer).

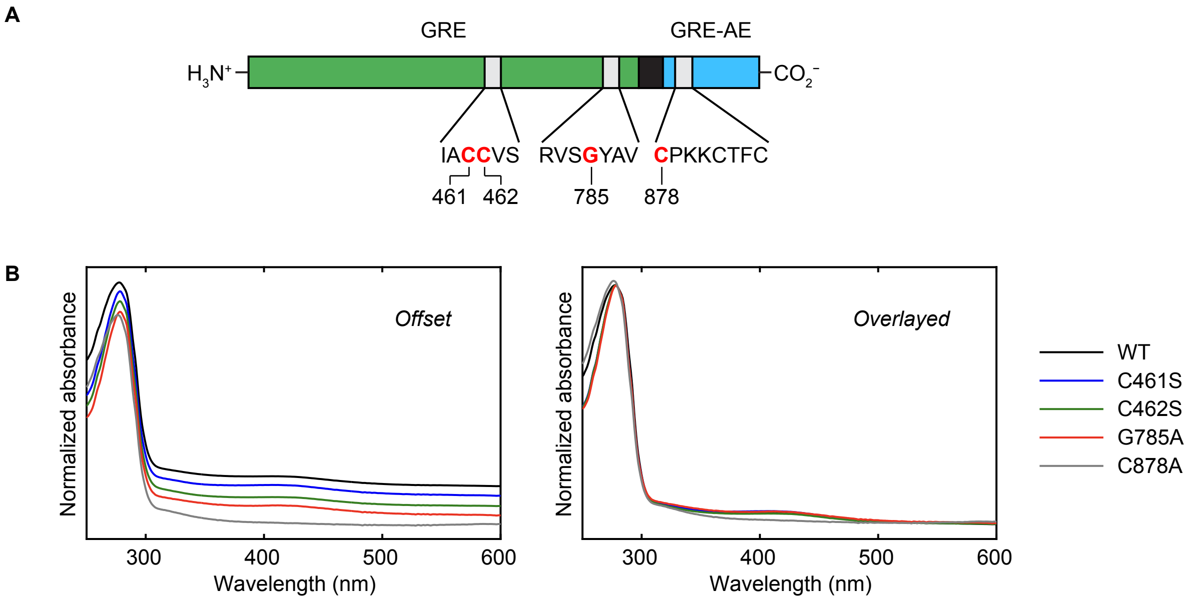


**Figure S4. (A)** Domain architecture of *Fs*GRE/GRE-AE. Conserved amino acid residues that were mutated are highlighted in red with their residue numbers shown below. **(B)** UV/vis spectra of *Fs*GRE/GRE-AE variants in as-isolated states. Identical set of spectra are displayed with an offset (*left*) or are overlayed with each other (*right*). Only the C878A variant is lacking the feature at ~410 nm, indicating the loss of an FeS cluster.

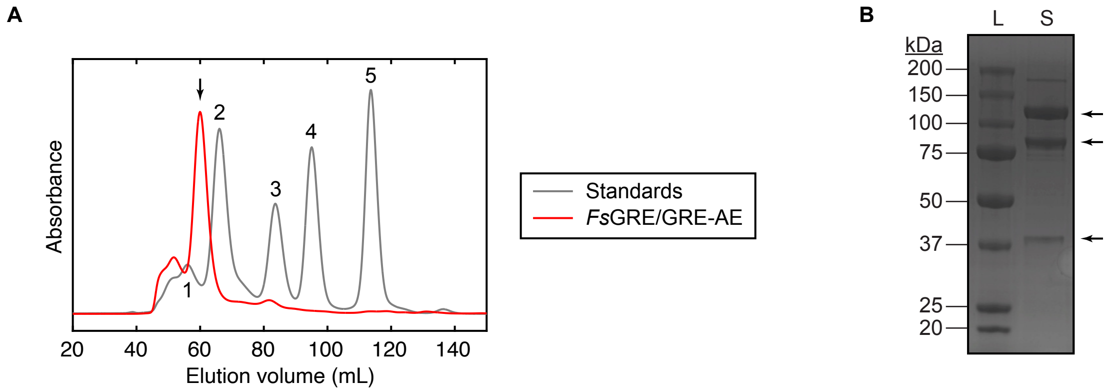


**Figure S5.** Anaerobic size-exclusion chromatography (SEC) of *Fs*GRE/GRE-AE. **(A)** The chromatogram of as-isolated *Fs*GRE/GRE-AE (red trace) overlayed with the chromatogram of a calibration standard mixture (grey trace). The major peak from the *Fs*GRE/GRE-AE injection is marked with an arrow. MW for protein standards: **1**, 440 kDa; **2**, 158 kDa; **3**, 67 kDa; **4**, 43 kDa; **5**, 15 kDa. **(B)** SDS-PAGE of the fraction from the major peak in SEC. L, protein ladder; S, sample. The same three bands observed in the as-isolated samples (see main text) are marked with arrows.

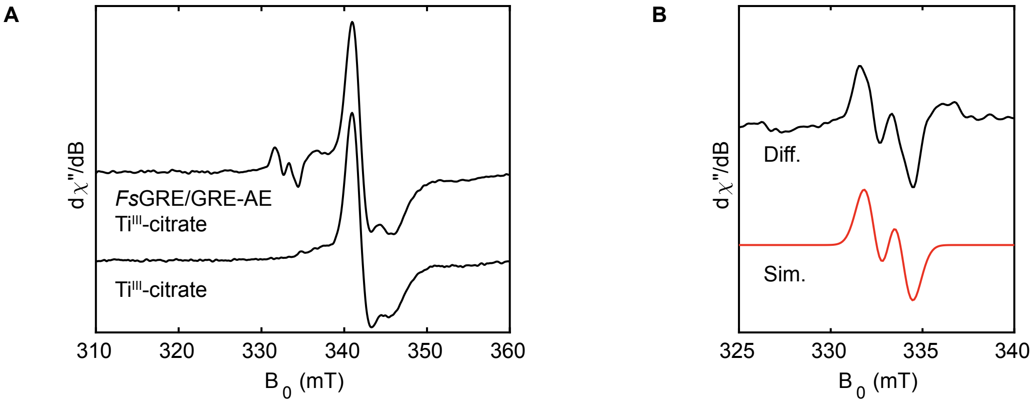


**Figure S6.** Activation of *Fs*GRE/GRE-AE using Ti$^{III}$-citrate. **(A)** X-band EPR spectrum of *Fs*GRE/GRE-AE activated with Ti$^{III}$-citrate, compared with the spectrum of Ti$^{III}$-citrate alone collected under the same conditions. **(B)** The difference spectrum yields the glycyl radical signal that matches well with the simulated spectrum in red. EPR conditions: temperature = 77 K, microwave frequency = 9.37 GHz, microwave power = 1 μW.

A

B

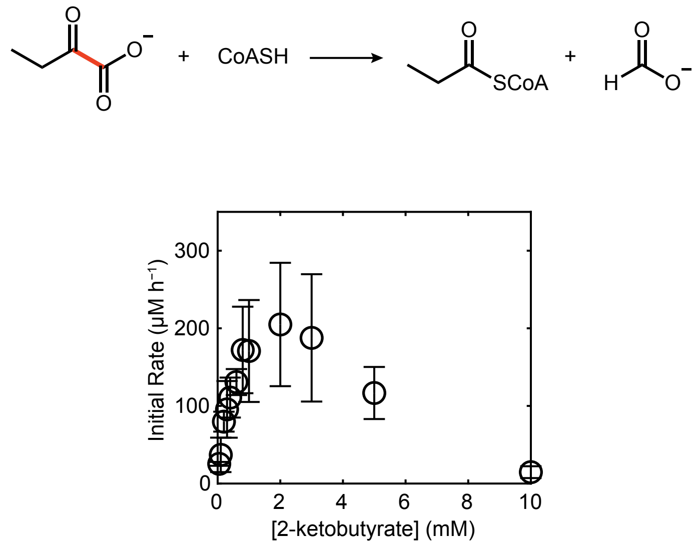


**Figure S7.** Activity assay of *Fs*GRE/GRE-AE with 2-ketobutyrate as the substrate. **(A)** Formate lyase reaction scheme for 2-ketobutyrate. **(B)** Enzyme kinetics under varying concentrations of 2-ketobutyrate as the substrate. Each data point with an error bar represents mean ± S.D. (standard deviation) of duplicate measurements. Assay conditions: [Enzyme] = 1 μM; [CoA] = 0.1 mM.

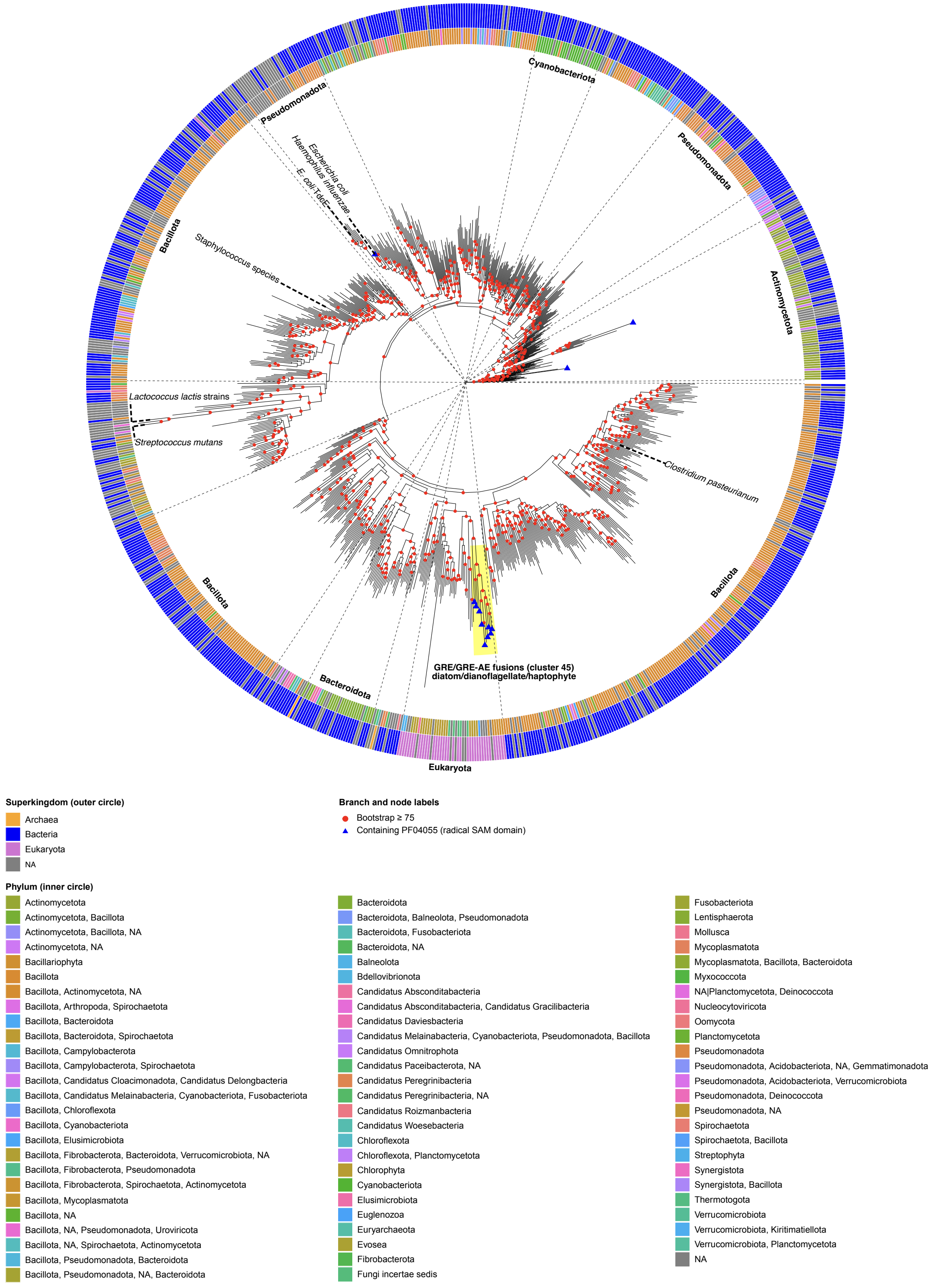


**Figure S8.** Maximum-likelihood phylogenetic tree of PFL sequences. The sequences of mega-cluster 1 from the AS 100 SSN were used to generate the tree. GRE/GRE-AE fusion proteins from cluster 45 (highlighted in yellow) form a distinct clade within the larger eukaryotic PFL clade.

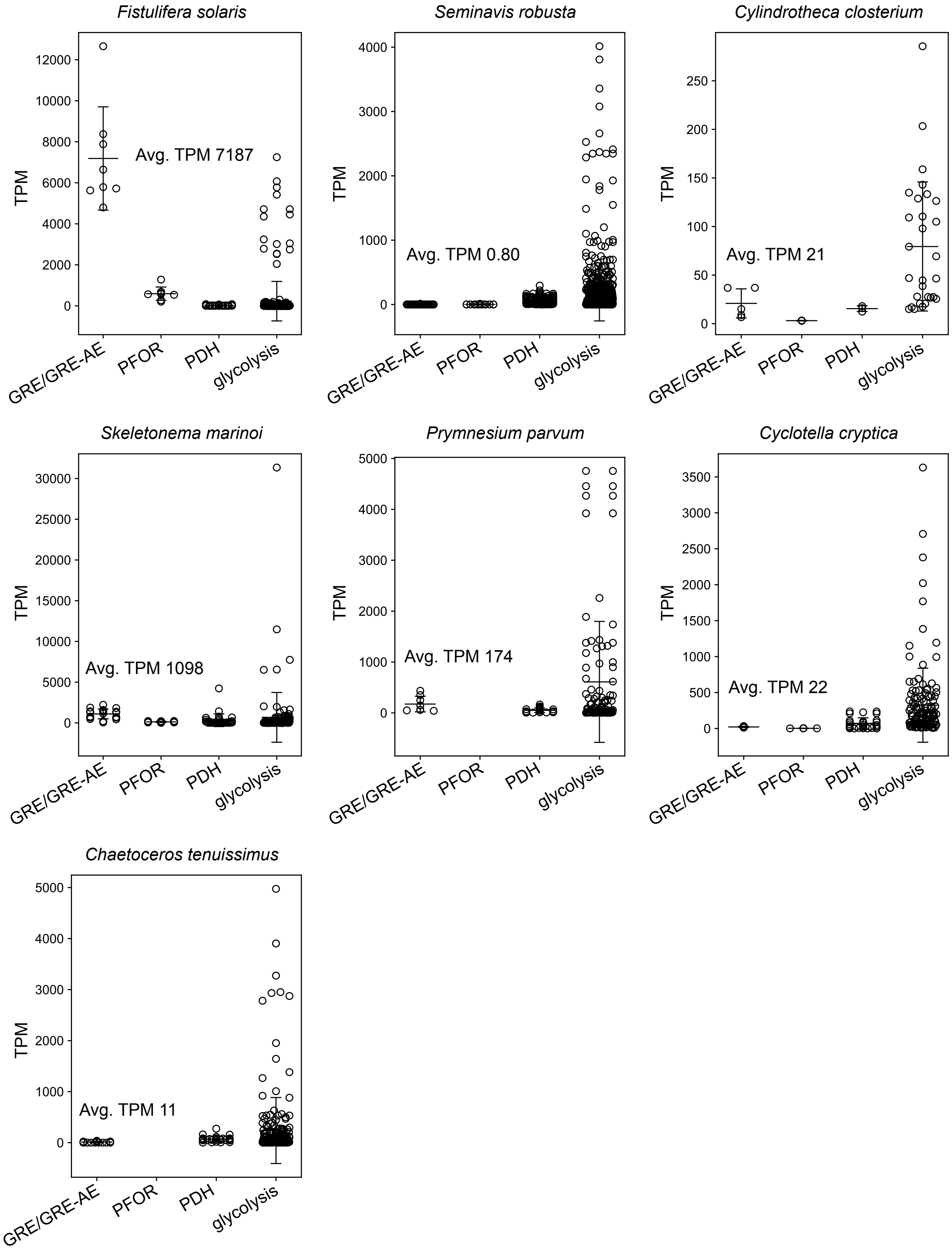


**Figure S9.** Transcriptomic analysis of organisms that encode GRE/GRE-AE fusion proteins. Each dot represents the gene expression level of a gene in the category from various biological samples. See **Supplementary Methods** for the analysis details.

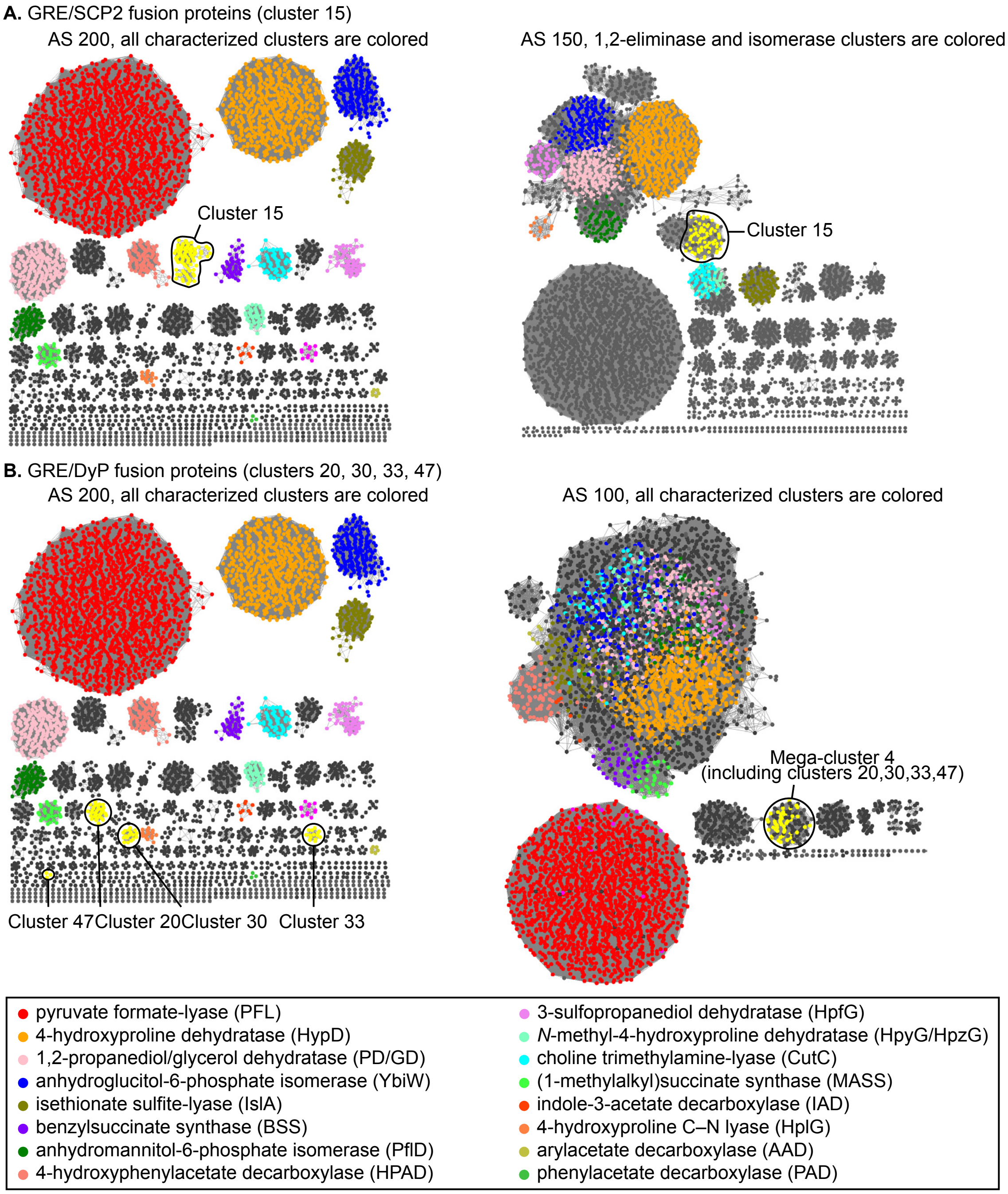


**Figure S10.** SSNs of PF02901-containing sequences at varying alignment score (AS) thresholds. Isofunctional clusters from the AS200 SSN are colored as described in figure. **(A)** Cluster 15, which contains GRE/SCP2 fusion protein sequences, is highlighted in yellow in the AS 200 SSN (*left*) and in the AS 150 SSN (*right*), showing its relationship to other 1,2-eliminases and isomerases. **(B)** Clusters 20, 30, 33, and 47, which contain GRE/DyP fusion proteins sequences, are highlighted in yellow in the AS 200 SSN (*left*) and in the AS 100 SSN (*right*). These clusters form the mega-cluster 4, which is distinct from all characterized GREs to date.

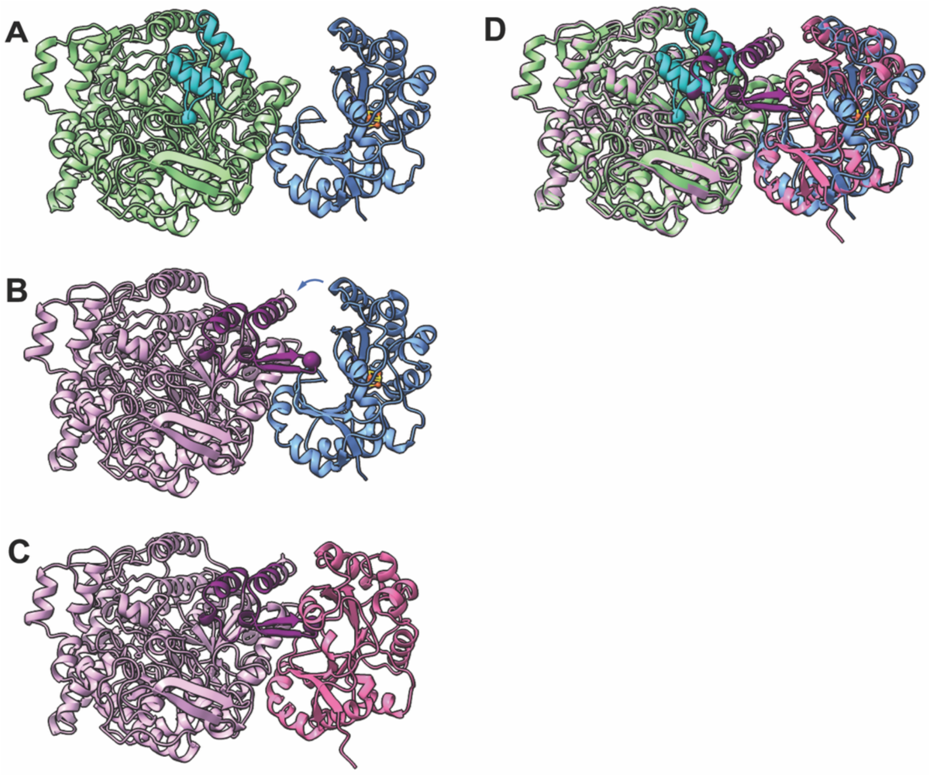


**Figure S11.** Orientations of *Ec*PFL with *Ec*PFL-AE from cryo-EM map and AlphaFold3 (AF3) modeling. **(A)** Orientation of *Ec*PFL protomer (PDB ID 1H18; gray) with *Ec*PFL-AE (PDB ID 3CB8; blue) from cryo-EM map. The “closed” GRD is highlighted in teal with the alpha carbon of G734 shown as a sphere. Sulfur atoms are shown in yellow and iron atoms in orange. **(B)** *Ec*PFL from AF3 modeling (light pink) with *Ec*PFL interacting with *Ec*PFL-AE (PDB ID 3CB8; blue) in the orientation from cryo-EM map. The “open” GRD is highlighted in dark purple with the alpha carbon of G734 shown as a sphere. The blue arrow indicates the movement of *Ec*PFL-AE in the orientation from cryo-EM map to achieve the position of *Ec*PFL-AE in AF3 modeling (shown in **C**) with *Ec*PFL. **(C)** AF3 modeling of *Ec*PFL with *Ec*PFL-AE. Same coloring for *Ec*PFL as in **B**. *Ec*PFL-AE is shown in hot pink. G734 is within the active site of PFL-AE. **(D)** Superposition of the ‘open GRD’ AF3 model (from panel **C**) aligned with ‘closed GRD’ structure of *Ec*PFL and *Ec*PFL-AE (from panel **A**). Same coloring from panels **A** and **C**.

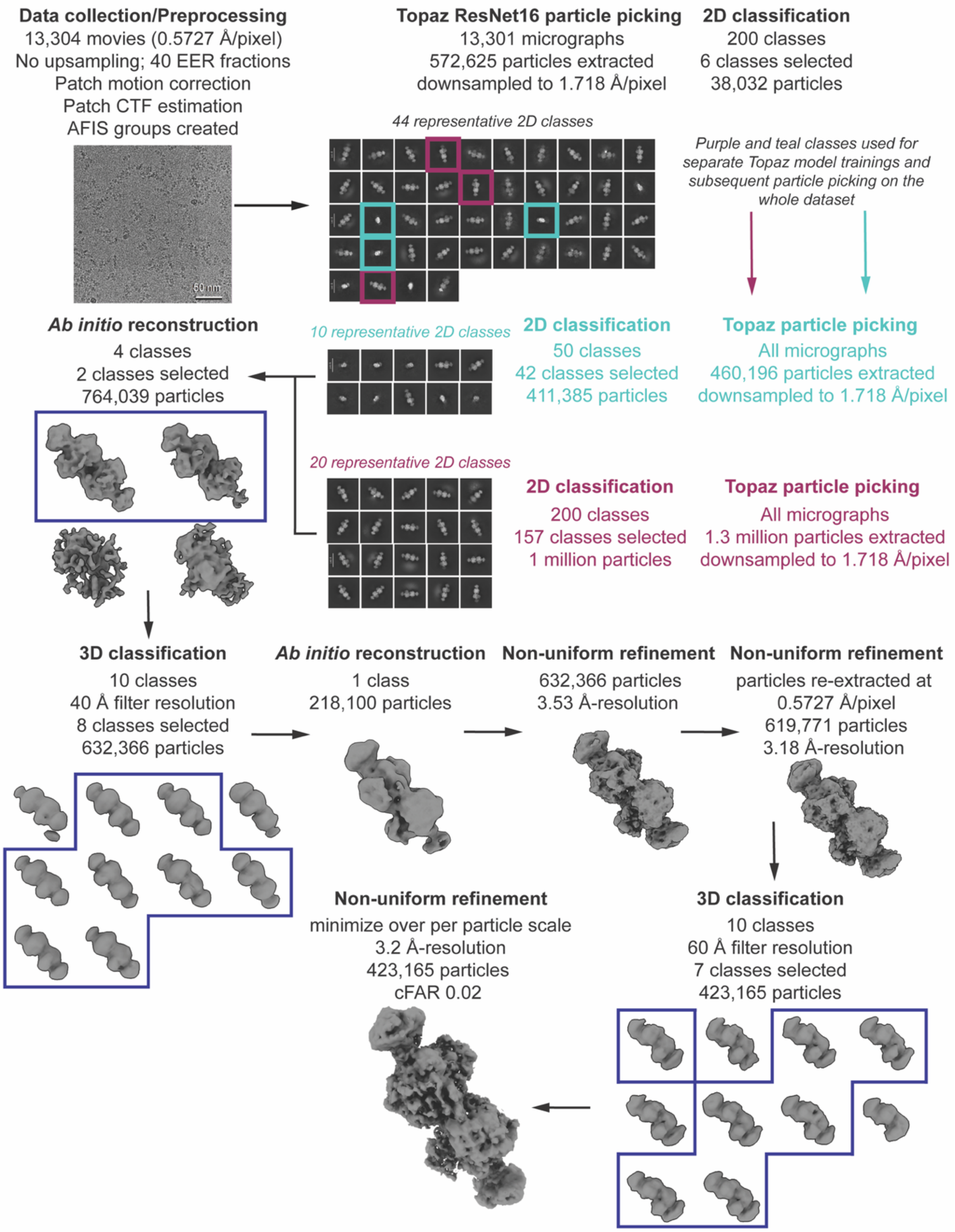


**Figure S12.** Cryo-EM data processing pipeline in cryoSPARC. Scheme describing the processing steps employed to arrive at the final 3.2-Å resolution *Fs*GRE/GRE-AE cryo-EM map. The final map, which unfortunately suffers from preferred orientation, is contoured to 5 sdLevel.

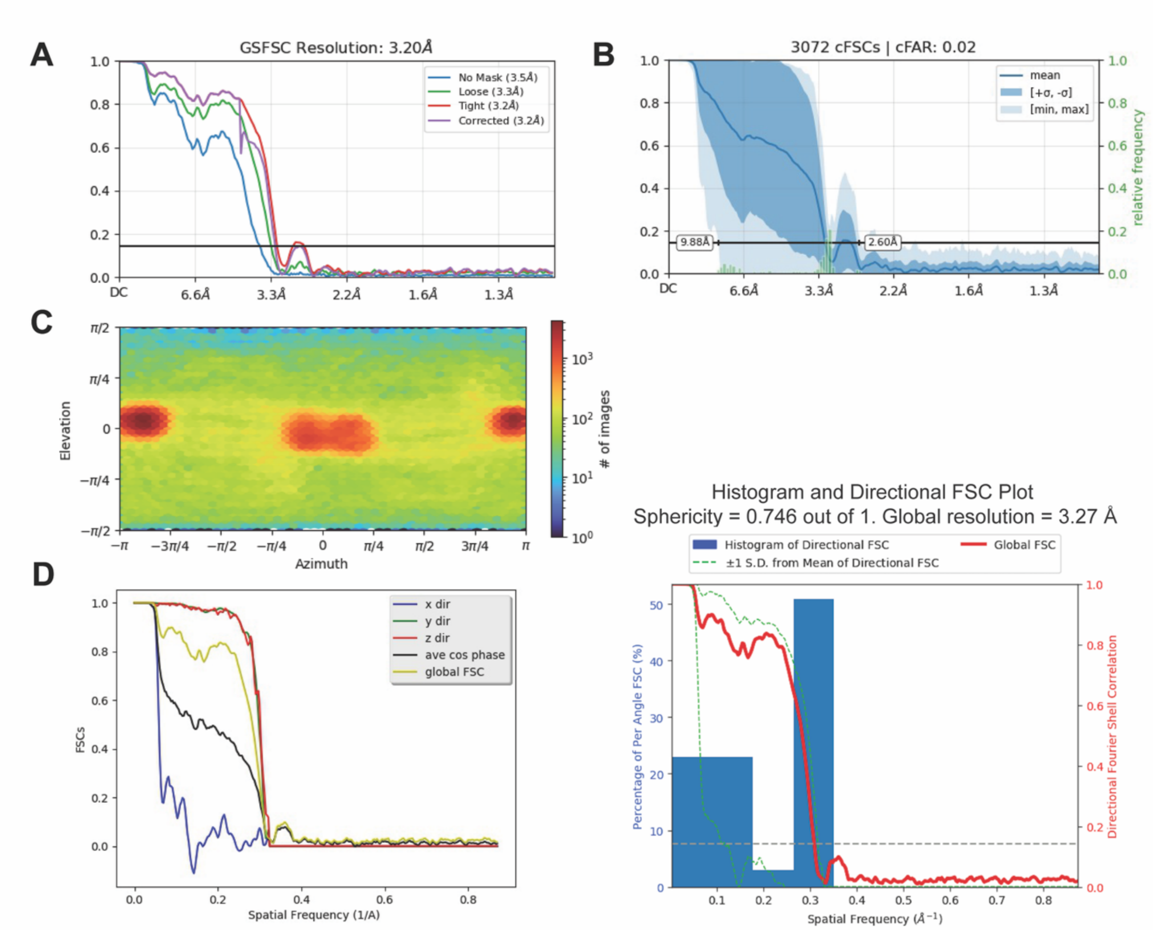


**Figure S13.** Fourier Shell Correlation (FSC) curves and angular distribution of the final *Fs*GRE/GRE-AE map. **(A)** Gold Standard FSC curve and final resolution as calculated by cryoSPARC. **(B)** Conical Fourier Shell Correlation (cFSC) plot from cryoSPARC. **(C)** Viewing direction distribution plot from cryoSPARC. **(D)** 3D FSC calculation using the Salk Institute for Biological Sciences remote 3D FSC server.

### C. Supplementary Tables

**Table S1.** Fe quantification of as-isolated *Fs*GRE/GRE-AE variants.

| **Variant** | **[Protein] (μM)**[a] | **[Total Fe] (mM)**[b] | **Fe per polypeptide**[c] |
|---|---|---|---|
| WT | 160 ± 10 | 0.70 ± 0.09 | 4.5 ± 0.6 |
| C461S | 100 ± 6 | 0.32 ± 0.09 | 3.2 ± 0.9 |
| C462S | 204 ± 7 | 0.92 ± 0.11 | 4.5 ± 0.6 |
| G785A | 96 ± 10 | 0.40 ± 0.09 | 4.2 ± 1.0 |
| C878A | 48 ± 5 | 0.016 ± 0.007 | 0.3 ± 0.1 |

[a] Estimated using BCA assay. Mean ± S.D. of triplicate measurements.
[b] Estimated using ferene assay. Mean ± S.D. of triplicate measurements.
[c] [Total Fe] ÷ [Protein] with standard error propagation.

**Table S2.** GRE/GRE-AE fusion proteins identified using iterative BLAST against the NCBI Non-redundant Protein Database.

| GenBank Accession | Organism | Taxonomy | Sequence Length | Partial Sequence |
|---|---|---|---|---|
| CEM20970.1 | *Chromera velia* CCMP2878 | alveolate | 1101 | |
| CAE0401902.1 | *Amphora coffeiformis* | diatom | 902 | |
| CAE0401901.1 | *Amphora coffeiformis* | diatom | 807 | |
| CAE0401904.1 | *Amphora coffeiformis* | diatom | 1032 | |
| CAE0401906.1 | *Amphora coffeiformis* | diatom | 1127 | |
| KAL3942348.1 | *Bacillariaceae sp.* | diatom | 1111 | |
| GAB7153730.1 | *Chaetoceros neogracilis* | diatom | 1173 | |
| GFH57376.1 | *Chaetoceros tenuissimus* | diatom | 1096 | |
| KAL7470471.1 | *Conticribra weissflogii* | diatom | 1181 | |
| CAD8336534.1 | *Craspedostauros australis* | diatom | 928 | |
| KAL3815393.1 | *Cyclostephanos tholiformis* | diatom | 1115 | |
| KAL3826620.1 | *Cyclostephanos tholiformis* | diatom | 1112 | |
| KAL3776605.1 | *Cyclotella atomus* | diatom | 1170 | |
| KAL7506221.1 | *Cyclotella atomus* | diatom | 1170 | |
| KAL7482200.1 | *Cyclotella cf. meneghiniana* | diatom | 994 | |
| KAL3788094.1 | *Cyclotella cryptica* | diatom | 1187 | y |
| CAJ1963009.1 | *Cylindrotheca closterium* | diatom | 1111 | |
| KAL3760044.1 | *Discostella pseudostelligera* | diatom | 1108 | |
| KAL3760034.1 | *Discostella pseudostelligera* | diatom | 1197 | |
| KAL7566397.1 | *Epithemia clementina (nom. ined.)* | diatom | 1104 | |
| GAX16364.1 | *Fistulifera solaris* | diatom | 1106 | |
| GAX28273.1 | *Fistulifera solaris* | diatom | 1107 | |
| KAL7453632.1 | *Mediolabrus comicus* | diatom | 1133 | |
| CAD8298895.1 | *Pseudictyota dubia* | diatom | 1156 | |
| CAD8298888.1 | *Pseudictyota dubia* | diatom | 1086 | |
| CAB9529030.1 | *Seminavis robusta* | diatom | 1086 | |
| CAB9515245.1 | *Seminavis robusta* | diatom | 969 | y |
| GAB5158302.1 | *Skeletonema ardens* | diatom | 1122 | y |
| GAB5182892.1 | *Skeletonema costatum* | diatom | 1152 | |
| GAB5206387.1 | *Skeletonema costatum* | diatom | 1152 | |
| GAB5236816.1 | *Skeletonema dohrnii* | diatom | 1133 | |
| GAB5246481.1 | *Skeletonema grevillei* | diatom | 1125 | y |
| GAB5262508.1 | *Skeletonema japonicum* | diatom | 1134 | y |
| KAK1739602.1 | *Skeletonema marinoi* | diatom | 1139 | |
| GAB5281421.1 | *Skeletonema menzelii* | diatom | 1118 | |
| KAL7493138.1 | *Skeletonema menzelii* | diatom | 1120 | |
| KAL7442423.1 | *Skeletonema potamos* | diatom | 739 | |
| GAB5289988.1 | *Skeletonema pseudocostatum* | diatom | 1137 | |
| GAB5309083.1 | *Skeletonema subsalsum* | diatom | 1086 | y |
| GAB5334583.1 | *Skeletonema tropicum* | diatom | 1133 | y |
| KAL3763716.1 | *Stephanodiscus triporus* | diatom | 1195 | |
| KAL9178626.1 | *Thalassiosira profunda* | diatom | 1142 | |
| KAL7543985.1 | *Thalassiosira sp.* AJA248-18 | diatom | 905 | y |
| CAE8605728.1 | *Polarella glacialis* | dinoflagellate | 923 | |
| CAE8641210.1 | *Polarella glacialis* | dinoflagellate | 934 | |
| XRB19841.1 | *Pseudoscourfieldia marina* | green algae | 1111 | |
| XRB05901.1 | *Pycnococcus provasolii* | green algae | 1115 | |
| KAL1508084.1 | *Prymnesium parvum* | haptophyte | 1079 | |
| KAL1522014.1 | *Prymnesium parvum* | haptophyte | 1079 | |

**Table S3.** SSN clusters containing ≥10 unique sequences that have average sequence lengths >950 amino acids.

| Cluster Number | Number of Sequences | Average Length | Additional Domain |
|---|---|---|---|
| 15 | 86 | 995 | SCP2 sterol binding |
| 20 | 56 | 1190 | DyP |
| 30 | 27 | 1311 | DyP |
| 33 | 24 | 1309 | DyP |
| 34 | 23 | 987 | - |
| 45 | 20 | 1105 | Radical SAM (GRE-AE) |
| 47 | 18 | 1136 | DyP |

### Table S4. Sequences of primers used for site-directed mutagenesis.

| Primer | Sequence (5′→3′) |
|---|---|
| C461S, N-fragment, forward | CAGCGGCCTGGTGCCGCGCGGCAGCCATATGATGTTTAAACCCAAGACAGTACTATCACA |
| C461S, N-fragment, reverse | CATGCCTGAAACACAGGATGCGATGCCGTAGTCGCAGCCG |
| C461S, C-fragment, forward | GACTACGGCATCGCATCCTGTGTTTCAGGCATGCGTATTGGTGTG |
| C461S, C-fragment, reverse | GATCTCAGTGGTGGTGGTGGTGGTGCTCGAGTTATGCAACGTTGCTCAGG |
| C462S, N-fragment, forward | *Identical to C461S, N-fragment, forward primer* |
| C462S, N-fragment, reverse | ACGCATGCCTGAAACAGAGCATGCGATGCCGTAGTCGCAG |
| C462S, C-fragment, forward | TACGGCATCGCATGCTCTGTTTCAGGCATGCGTATTGGTG |
| C462S, C-fragment, reverse | *Identical to C461S, C-fragment, reverse primer* |
| G785A, N-fragment, forward | *Identical to C461S, N-fragment, forward primer* |
| G785A, N-fragment, reverse | ACGCACGGCATAGGCGCTAACACGAATGGTCAGGTTCGGG |
| G785A, C-fragment, forward | ACCATTCGTGTTAGCGCCTATGCCGTGCGTTTTTGTCGTC |
| G785A, C-fragment, reverse | *Identical to C461S, C-fragment, reverse primer* |
| C878A, N-fragment, forward | *Identical to C461S, N-fragment, forward primer* |
| C878A, N-fragment, reverse | GCACTTTTTCGGGGCGCCCTGCAAGAAAACGTTGG |
| C878A, C-fragment, forward | GTTTTCTTGCAGGGCGCCCCGAAAAAGTGCACCTT |
| C878A, C-fragment, reverse | *Identical to C461S, C-fragment, reverse primer* |

**Table S5.** Cryo-EM data collection

| | |
|---|---|
| Microscope | FEI Titan Krios (PNCC Krios-1) |
| Camera | Falcon 4i DED |
| Acceleration voltage (kV) | 300 |
| Spherical aberration (mm) | 2.7 |
| Magnification (x) | 215,000 |
| Pixel size (Å) | 0.5727 |
| Defocus range (µm) | −0.5 to −2.0 |
| Exposure time (s) | 3.63 |
| Total exposure ($e^-$/Å$^2$) | 50 |
| Shots per hole | 7 |
| Total movies collected | 13,304 |
| Collection software | EPU |
